# Reprogramming neuroblastoma by diet-enhanced polyamine depletion

**DOI:** 10.1101/2024.01.07.573662

**Authors:** Sarah Cherkaoui, Lifeng Yang, Matthew McBride, Christina S. Turn, Wenyun Lu, Caroline Eigenmann, George E. Allen, Olesya O. Panasenko, Lu Zhang, Annette Vu, Kangning Liu, Yimei Li, Om H. Gandhi, Lea Surrey, Michael Wierer, Eileen White, Joshua D. Rabinowitz, Michael D. Hogarty, Raphael J. Morscher

**Affiliations:** Pediatric Cancer Metabolism Laboratory, Children’s Research Center, University of Zurich, 8032 Zurich, Switzerland; Division of Oncology, University Children’s Hospital Zurich and Children’s Research Center, University of Zurich, 8032 Zurich, Switzerland; Department of Chemistry, Princeton University, Princeton, NJ 08544, USA; Ludwig Institute for Cancer Research, Princeton Branch, Princeton University, Princeton, NJ 08544, USA; Division of Oncology and Department of Pediatrics, Children’s Hospital of Philadelphia, Philadelphia, PA 19104, USA; Perelman School of Medicine at the University of Pennsylvania, Philadelphia, PA 19104, USA; Bioinformatics Support Platform, Faculty of Medicine, University of Geneva 1211, Switzerland; Department of Microbiology and Molecular Medicine, Institute of Genetics and Genomics Geneva, Faculty of Medicine, University of Geneva, 1211 Geneva, Switzerland; BioCode: RNA to proteins (R2P) Platform, University of Geneva, 1211 Geneva, Switzerland; Department of Molecular Biology and Biochemistry, Rutgers University, Piscataway, NJ 08901, USA; Department of Molecular Biology and Biochemistry, Rutgers Cancer Institute of New Jersey, New Brunswick, NJ 08901, USA; Department of Pathology and Laboratory Medicine, Children’s Hospital of Philadelphia, Philadelphia, PA 19104, USA; Proteomics Research Infrastructure, Panum Institute, Blegdamsvej 3B, University of Copenhagen, 2200 Copenhagen, Denmark; Division of Human Genetics, Medical University Innsbruck, Peter-Mayr-Str. 1, 6020 Innsbruck, Austria

**Keywords:** Protein translation, polyamines, difluoromethylornithine, amino acid restriction, diet, proline, arginine, glutamine, ornithine, ornithine aminotransferase

## Abstract

Neuroblastoma is a highly lethal childhood tumor derived from differentiation-arrested neural crest cells^1,2^. Like all cancers, its growth is fueled by metabolites obtained from either circulation or local biosynthesis^3,4^. Neuroblastomas depend on local polyamine biosynthesis, with the inhibitor difluoromethylornithine showing clinical activity^5^. Here we show that such inhibition can be augmented by dietary restriction of upstream amino acid substrates, leading to disruption of oncogenic protein translation, tumor differentiation, and profound survival gains in the TH-*MYCN* mouse model. Specifically, an arginine/proline-free diet decreases the polyamine precursor ornithine and augments tumor polyamine depletion by difluoromethylornithine. This polyamine depletion causes ribosome stalling, unexpectedly specifically at adenosine-ending codons. Such codons are selectively enriched in cell cycle genes and low in neuronal differentiation genes. Thus, impaired translation of these codons, induced by the diet-drug combination, favors a pro-differentiation proteome. These results suggest that the genes of specific cellular programs have evolved hallmark codon usage preferences that enable coherent translational rewiring in response to metabolic stresses, and that this process can be targeted to activate differentiation of pediatric cancers.

**Graphical Abstract:** 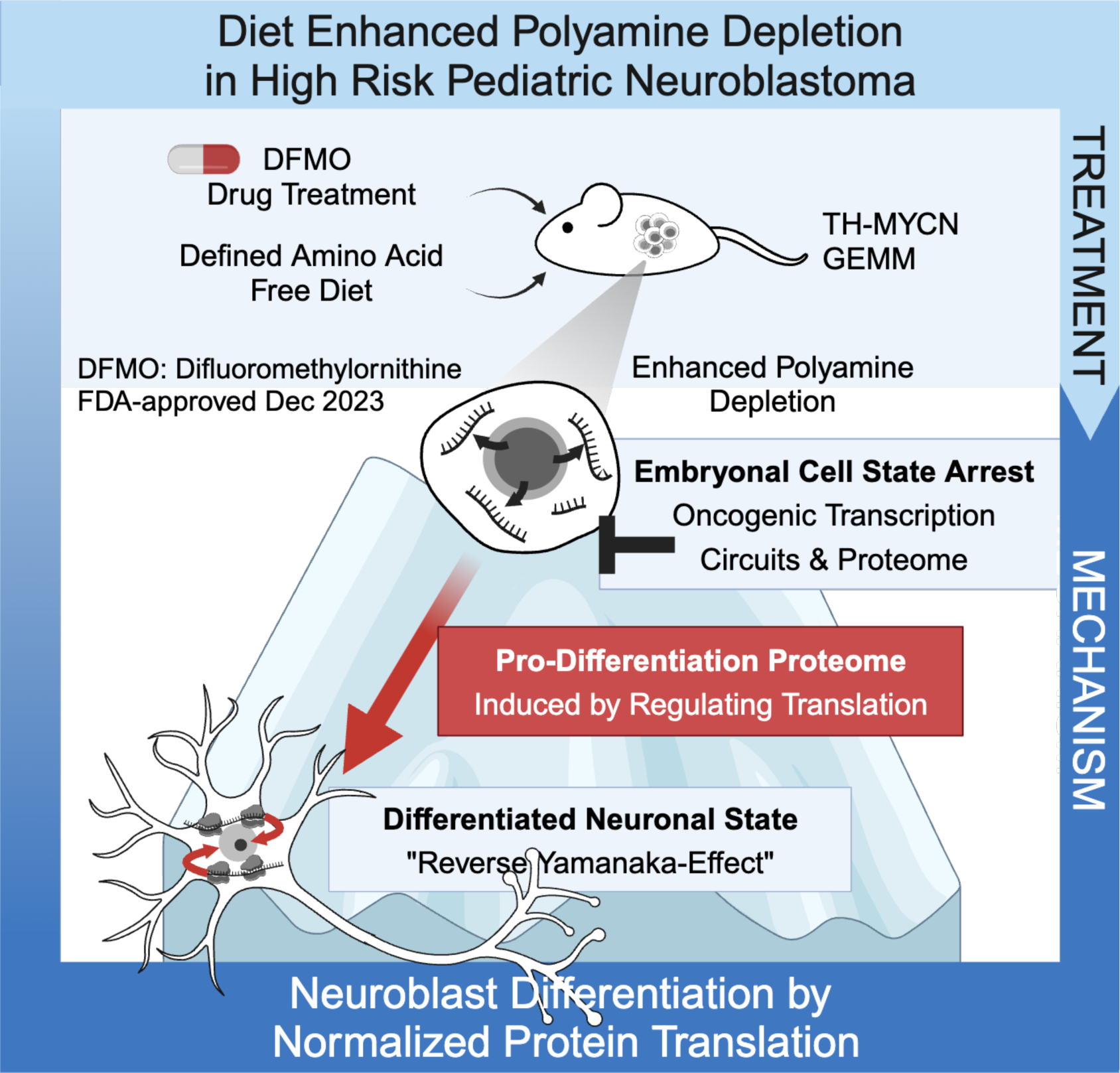

**Highlights:**

- Extra-tumoral conversion of arginine feeds tumor ornithine via uptake from circulation in MYCN-neuroblastoma.
- A proline and arginine free diet enhances pharmacological polyamine depletion via reduced ornithine substrate availability.
- Polyamine depletion disrupts oncogenic translation to induce a pro-differentiation proteome causing neuroblast differentiation and prolonged survival in the TH-MYCN mouse model.
- Genes of specific cellular programs have evolved codon usage preferences that enable coherent translational rewiring in response to metabolic stress, such as polyamine depletion.

## Introduction

Hyperactive MYC signaling via amplification of the *MYCN* proto-oncogene is a hallmark of neuroblastoma^6^ and drives aggressive disease with poor outcome^7–9^. In the TH-*MYCN* genetically engineered mouse model enforced *MYCN* expression in sympathoadrenal cells induces neuroblastomas^10^. The rate-limiting enzyme in polyamine biosynthesis, ornithine decarboxylase (ODC), is directly transcriptionally upregulated by MYC(N)^11^. Inhibition of ODC via difluoromethylornithine (DFMO) has recently received pre-approval by the FDA’s Oncologic Drug Advisory Committee (ODAC) for prevention of relapse in children with high-risk neuroblastoma^5^. Combined treatment strategies to enhance DFMO activity are therefore of great interest.

One potential such strategy is depletion of the ODC substrate ornithine. Ornithine can be derived from arginine via a single enzymatic step catalyzed by arginase, or from proline and glutamine via two or three steps converging on the enzyme ornithine amino transferase (OAT)^12^. In early childhood, most ornithine comes from proline via OAT^13,14^. Since neuroblastomas arise in early childhood, they might similarly depend on OAT, which was recently implicated as a metabolic driver in pancreatic cancer through the provision of ornithine and thereby polyamines^15^.

## Proline and arginine metabolism is altered in *MYCN*-driven neuroblastoma

We found that proline, whose catabolism can feed into ornithine via OAT, is strongly upregulated in MYCN-driven neuroblastoma. This was shown in three contexts: *MYCN*-amplified primary patient tumors (relative to unamplified tumors), xenografts induced from high *MYCN*-expressing patient-derived neuroblastoma cell lines (relative to low-expressing cell lines), and TH-*MYCN* genetically engineered murine tumors (relative to normal organs of the mouse) (Fig. 1a-d, Extended Data Fig1, Supplementary Table S1). In the TH-MYCN model, proline was also markedly higher in late tumors (> 50 mm^3^) compared to early tumors (Extended Data Fig.1h). Neither ornithine nor its other upstream precursors were consistently elevated. Thus, we identified proline as a strongly elevated metabolite in MYCN-driven neuroblastoma and a candidate to restrict for potential enhancement of neuroblastoma therapy.

**Figure 1:**
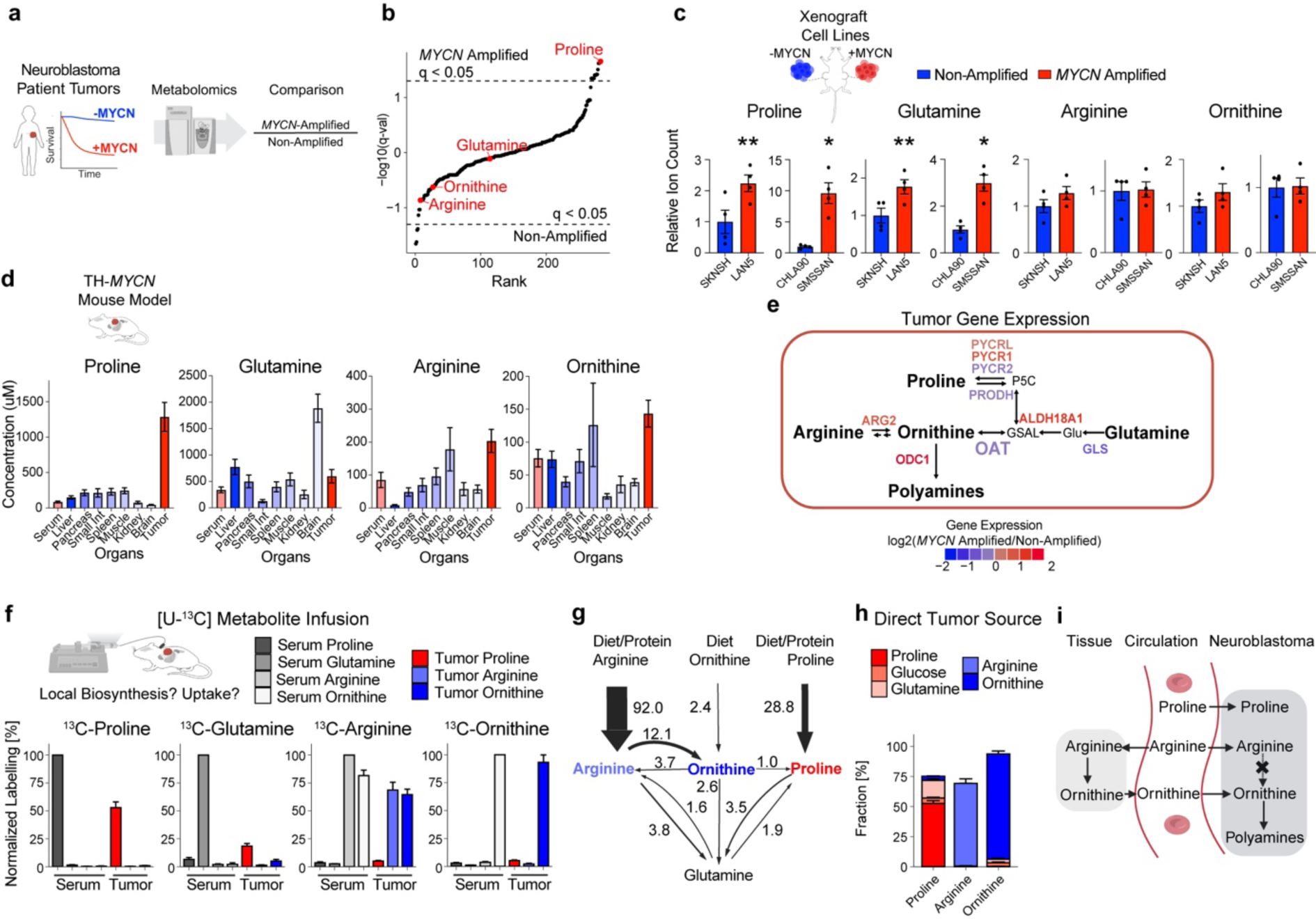
*MYCN*-driven neuroblastoma tumors are characterized by high proline and a functionally disconnected proline and arginine metabolism dependent on uptake from circulation. a) Primary neuroblastoma tumor tissue undergoing liquid chromatography-mass spectrometry-based metabolomics. b) Differential abundance of 303 metabolites. Proline is the most significantly increased metabolite in *MYCN* amplified primary human neuroblastoma relative to non-amplified tumors. Dotted line marks significance threshold, with p-values corrected for false discovery rate of 0.05, *q < 0.05. *n* = 10. c) Relative levels of proline, glutamine, arginine and ornithine in contralateral xenografts from *MYCN* amplified and non-amplified neuroblastoma cell lines. *P < 0.05, **P < 0.01, two-tailed paired t-test. Mean ± s.e.m., *n* = 4. d) Proline concentration in *MYCN* driven neuroblastoma tumors is significantly increased in the TH-*MYCN* mouse model^10^, whereas glutamine, arginine and ornithine levels across organs are within physiological range. Mean ± s.e.m., tumor proline *n* = 31, glutamine *n* = 29, arginine *n* = 27 and ornithine *n* = 24; other organs *n* = 8-31. e) Gene expression of the metabolic network producing the polyamine precursor ornithine highlights low OAT (ornithine amino transferase). Color of labels indicates relative expression in patients (*MYCN* amplified *n* = 93 / non-amplified *n* = 551)^21^. f) *In vivo* stable isotope tracing elucidates the precursors of intratumoral metabolites. Labelling is given normalized to the serum of each infused [U-^13^C]metabolite in TH-*MYCN* mice. Mean ± s.e.m., *n* = 4-9. g) Whole-body flux model^22^ of sources and interconversions between circulating metabolites. Exchange fluxes are given for circulating proline, ornithine and arginine and their exchange with glutamine. Flux in nmol C/min/g. Mean*, n* = 4-9. h) Direct circulating nutrient contributions to tumor tissue metabolite pools of proline, arginine and ornithine in TH-*MYCN* mice. The color indicates the respective circulating nutrient source. Mean ± s.e.m., *n* = 4-9. i) Schematic showing tumor metabolite sources in neuroblastoma. The non-essential amino acids proline and arginine are primarily taken up from circulation. Tracing identifies the polyamine precursor ornithine to be primarily derived from circulation and not from intratumoral biosynthesis from either arginine or from proline/glutamine through OAT.

As a complementary approach to identify potential metabolic targets for enhancing neuroblastoma therapy, we investigated the *in vivo* sources of proline and ornithine using stable isotope tracing in the TH-*MYCN* model^16–18^. Tumor proline came substantially both from circulating proline and from glutamine, reflecting the tumor acquiring its high levels from a combination of circulatory uptake and de novo biosynthesis. Unlike previous studies in newborns^19,20^ or pancreatic cancer^15^, this proline was not converted to ornithine in substantial quantities in neuroblastoma. Examination of primary tumor gene expression data revealed that OAT is actually low in MYCN-driven neuroblastoma, consistent with the need for an alternative ornithine source rather than synthesis from glutamine or proline. Isotope tracing confirmed the source to be circulating arginine and ornithine itself. Most circulating ornithine came from arginine. Tumor ornithine was most strongly labeled from circulating ornithine, although it was also strongly labeled from circulating arginine (Fig. 1e-f, Extended Data Fig.2). Quantitative modelling analysis of tumor ornithine sources revealed that arginine feeds tumor ornithine mainly indirectly, after being converted to ornithine elsewhere in the body and the resulting circulating ornithine is taken up by the tumor (Fig.1g-i, Extended Data Fig.3). Nevertheless, the ultimate upstream source of most neuroblastoma ornithine is arginine, highlighting arginine restriction as a potential complement to proline restriction and DMFO for neuroblastoma therapy.

## Combination of a proline and arginine-free diet with DMFO treats murine neuroblastoma

Using the TH-MYCN model, we examined the impact of combined dietary amino acid depletion (proline and arginine, Supplementary Table S2) with or without pharmacological ODC inhibition by DMFO (Fig.2a). Mice fed the proline and arginine free diet alone (ProArg-free) showed reduced neuroblastoma growth compared to control diet (CD), but without an effect on tumor free survival. As with prior studies, inhibition of polyamine biosynthesis by DFMO monotherapy extended survival^23–25^. For the mice with prolonged survival lethal tumor progression was observed after treatment cessation.

**Figure 2:**
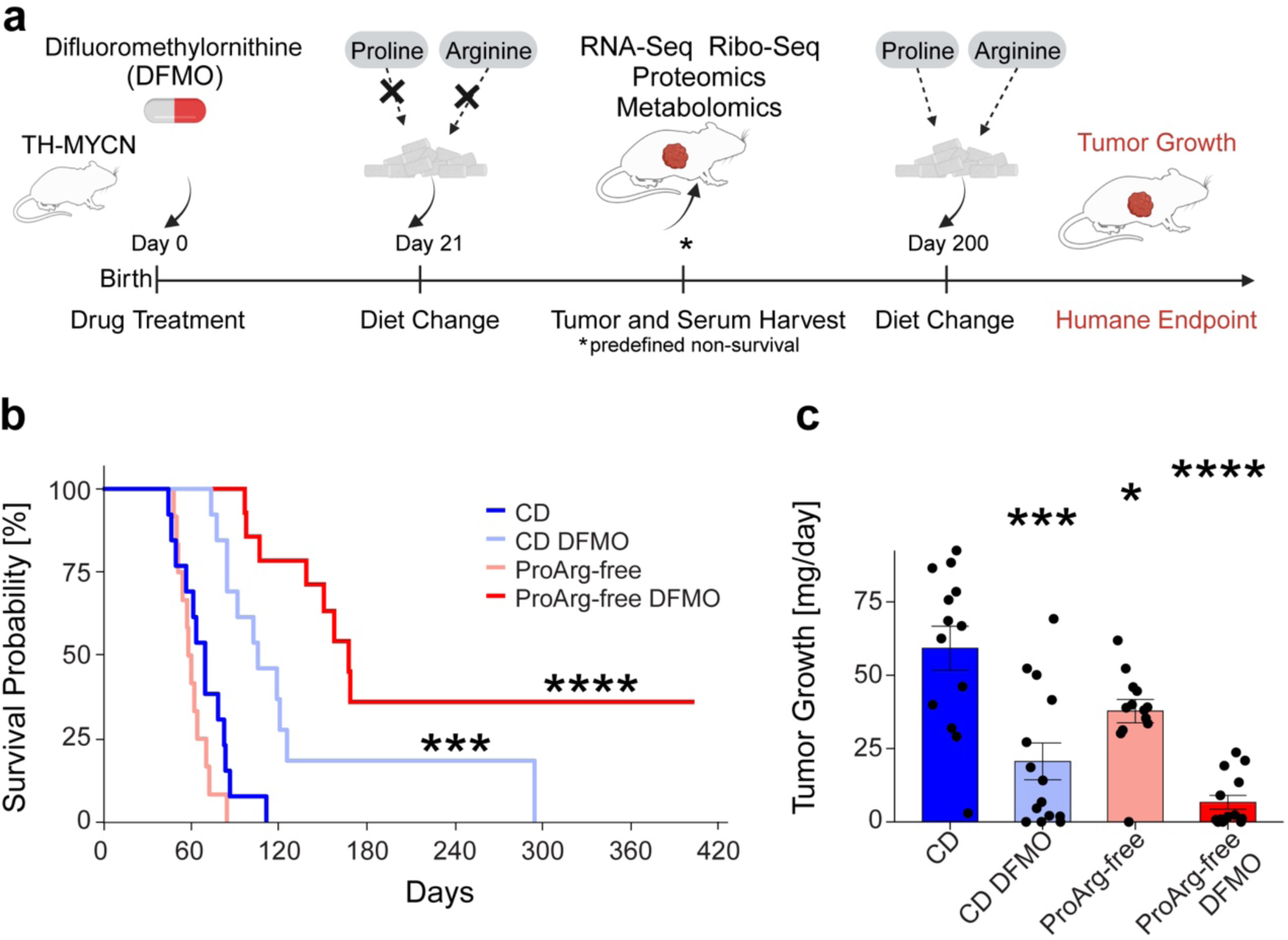
A proline/arginine free diet enhances tumor growth suppression by DFMO in *MYCN*-driven neuroblastoma. a) Schematic of 2 factor intervention including a combined proline and arginine amino acid free diet (day 21) and DFMO drug treatment via the drinking water (1 %, day 0) in the TH-*MYCN* genetically modified mouse model^10^. b) Kaplan-Meier curve of tumor free survival upon treatment where control diet (CD) or a proline and arginine free diet (ProArg-free) are combined with DFMO (difluoromethylornithine). Log-rank test p-value compared to CD. c) Tumor growth defined as tumor weight at death normalized by day of life. Two-tailed t-test compared to CD. Mean ± s.e.m.. For b and c: *P < 0.05, **P < 0.01, ***P < 0.001, ****P < 0.0001. CD *n* = 13, CD DFMO *n* = 14, ProArg-free *n* = 13, ProArg-free DFMO *n* = 14.

Strikingly, combining dietary proline and arginine removal with DFMO induced a marked survival benefit (Fig.2b), decreased tumor growth (Fig.2c), and increased time to detectable tumor (Extended Data Fig.4a-b). A third of mice in the ProArg-free DFMO regimen had extended survival, with ∼ 20 % remaining tumor-free as confirmed via necropsy. While the ProArg-free diet caused a reduction in mouse weight, this did not affect survival and was not worsened by adding DFMO (Extended Data Fig.4c). In summary, the ODC inhibitor DFMO, recently pre-approved by ODAC for neuroblastoma treatment, combined with a diet free of the non-essential amino acids arginine and proline, significantly augments anti-tumor activity with roughly one-fifth of treated mice remaining tumor free 100 days beyond the end of therapy.

## Arginine and proline-free diet depletes circulating ornithine and, together with DMFO, tumor polyamines

To reveal the metabolic reprogramming underlying this anti-tumor activity, we performed serum metabolomics (Fig.3a and Extended Data Fig.5). Across the entire metabolome, the metabolite showing the most significant decrease in serum in response to the ProArg-free diet was ornithine, the crucial polyamine precursor that we sought to deplete. The next two most significant metabolites were proline and arginine themselves (Fig.3b-c and Extended Data Fig.5b-c). Other notably altered metabolites included increased glutamine and decreased citrulline and the collagen breakdown product hydroxyproline (Extended Data Fig.5c). Arginine, proline, and ornithine were also significantly decreased in tumors, but to a lesser extent than in serum (Fig.3d and Extended Data Fig.5d-e). The depletion of intratumoral ornithine manifested despite raised tumor glutamine that gives rise to ornithine in other cancers via OAT^15^.

**Figure 3:**
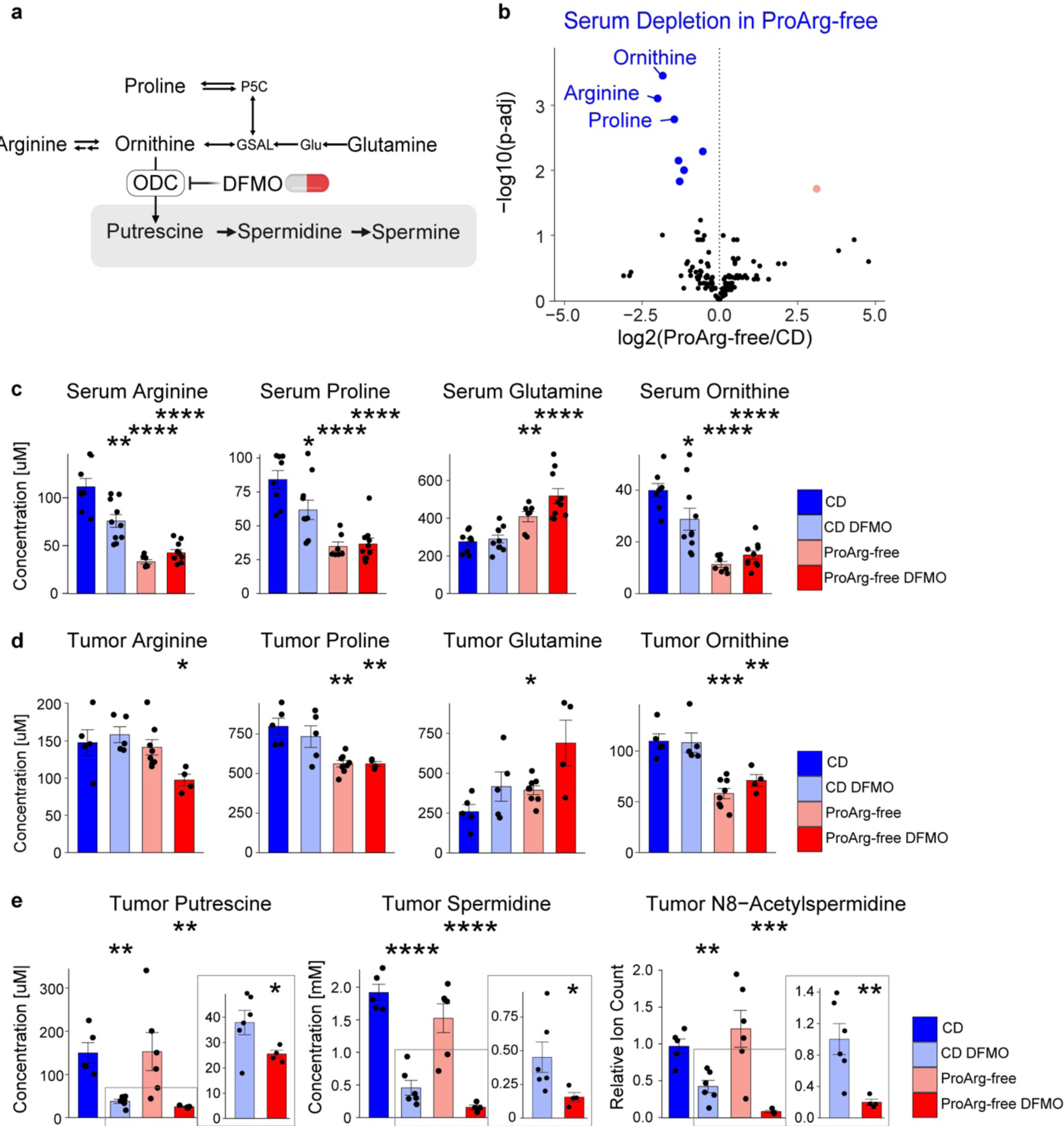
Dietary intervention causes substrate depletion to enhance polyamine biosynthesis inhibition by DFMO. a) Schematic of arginine and proline metabolism and its direct link to polyamines via ornithine. b) Differential serum metabolite levels comparing ProArg-free diet to CD. Blue highlights metabolites that are significantly depleted (FDR < 0.05) and rose upregulated compared to CD. *n* = 7-8. c) Serum arginine, proline, glutamine and ornithine across groups. Statistics comparisons to CD. Mean ± s.e.m., *n* = 7-10. d) Tumor arginine, proline, glutamine and ornithine levels reveals dysregulation of arginine and proline metabolism under combined treatment. Average age at sacrifice is 8 weeks. Statistics comparisons to CD. Mean ± s.e.m., *n* = 4-7. e) A ProArg-free diet enhances polyamine depletion in tumor tissue induced by DFMO treatment in prolonged treatment. Average age sacrifice 12 weeks. Comparison treatment to CD. Zooms display differences in polyamine levels between induced by ProArg-free on top of DFMO. Mean ± s.e.m., *n* = 4-6. *P < 0.05, **P < 0.01, ***P < 0.001, ****P < 0.0001, two-tailed t-test. Abbreviations: CD, control diet; ProArg-free, proline arginine free diet; DFMO, difluoromethylornithine.

Targeted LC-MS/MS measurements of tumor polyamines revealed that DMFO treatment decreased putrescine, the direct product of ornithine decarboxylation by ODC, and its derivatives such as spermidine. The ProArg-free diet potentiated the DFMO effect to further decrease polyamine levels, achieving a >10-fold reduction in spermidine compared to control diet and a >2-fold compared to DFMO monotherapy (Fig.3e). N8-acetylspermidine and N-acetyl-putrescine were also decreased by the diet-drug combination consistent with reduced catabolic flux (Fig.3e and Extended Data Fig.5f). Thus, dual dietary amino acid restriction depletes the key polyamine precursor ornithine and, in concert with DMFO, leads to enhanced tumor polyamine depletion.

## Diet-drug combination results in impaired eIF5A hypusination and translation defects

Polyamines stimulate translation and cell growth^26^. Arginine and proline also directly feed into translation as proteinogenic amino acids. To investigate the impact of the ProArg-free diet and DMFO on translation, we carried out ribosome profiling^27^ (Fig.4a and, quality control Extended Data Fig.6). This approach allows to identify globally which transcripts are loaded with ribosomes, and at which codons the ribosomes sit. Increased ribosome density can indicate sites of stalled translation, either due to uncharged tRNAs or a translation-machinery defect. Upon the ProArg-free diet, DFMO or the combination, ribosome occupancy was shifted towards the start codon and decreased at early elongation of the protein-encoding transcript (insert). Exclusively under combined diet-drug treatment, ribosomes accumulated at stop codons indicating defective ribosome release (Fig.4b). Such elongation and termination defects have been reported in cell models functionally deficient in eIF5A^28^, a translation factor post-translationally modified by the polyamine spermidine^29–31^. As this suggested *in vivo* eIF5A dysfunction, we probed eIF5A hypusination status (i.e., spermidine modification of eIF5A) across treatment groups. Whereas all tumors from the CD and ProArg-free groups without drug demonstrated complete eIF5A hypusination, reflecting sufficient spermidine for this purpose, incompletely hypusinated eIF5A was detected in one tumor treated with DFMO alone and a majority of the ProArg-free DFMO group (Extended Data Fig.7a-c).

**Figure 4:**
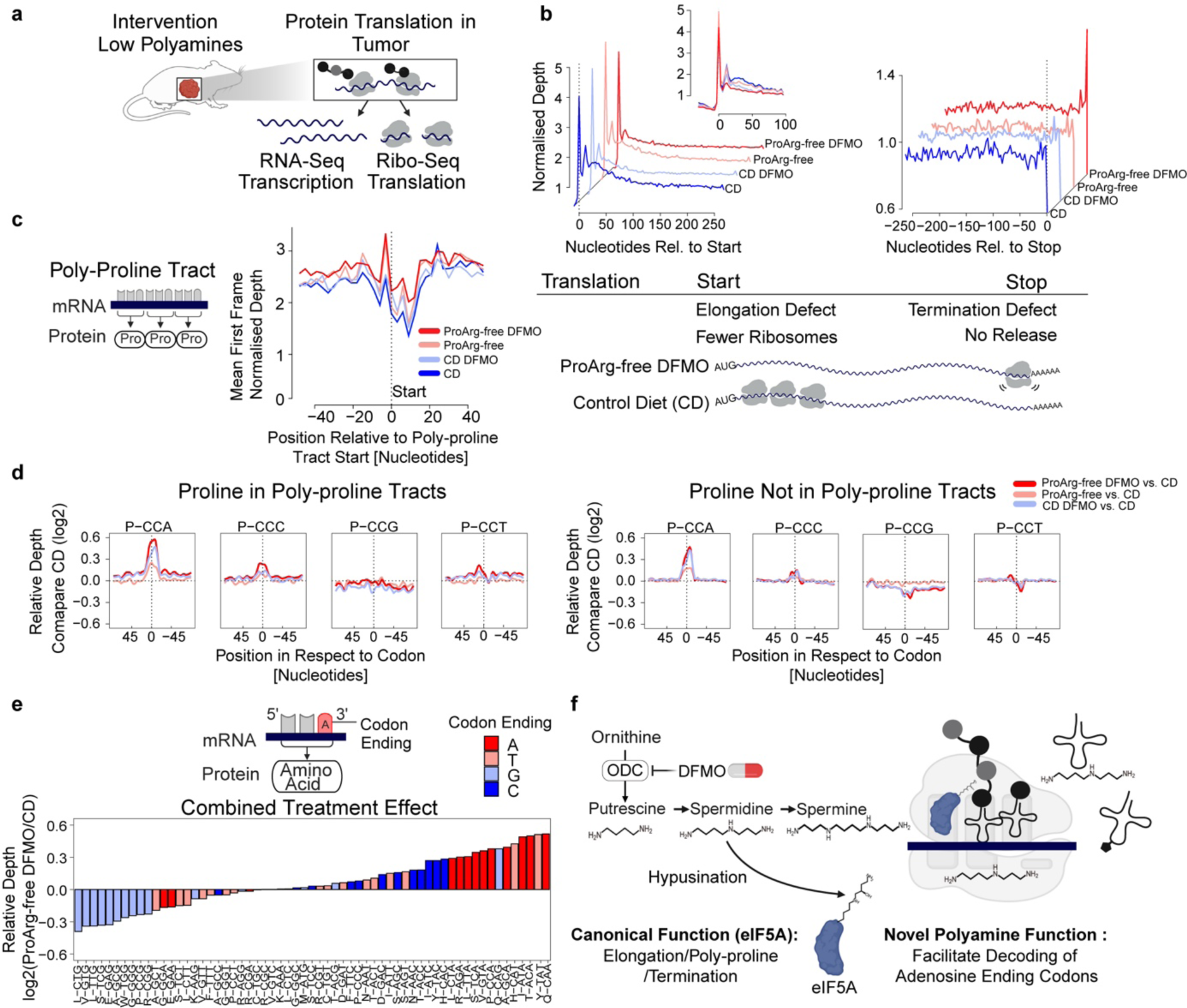
Ribo-Seq reveals defective decoding of adenosine-ending codons upon polyamine depletion. a) For functional evaluation of translation, tumors were lysed under the presence of a translation inhibitor for preparation of RNA- and Ribo-Seq libraries. Ribosome protected RNA fragments were isolated and sequenced for translation evaluation. b) Normalized ribosome density across transcripts. Combining a proline/arginine free diet with DFMO affects the global ribosome distribution at initiation/early elongation (left + insert) and causes a termination defect (right). c) Normalized ribosome depth at positions encoding for >= 3 prolines in a row. Decoding of these polyproline tracts is affected by combining DFMO with proline and arginine free diet. d) Proline translation defects are codon specific. Relative ribosome density centered around proline codons across treatment groups relative to control diet (zero-line). The left panels denote density of ribosomes at poly proline tracts and right codon occupancy on proline codons outside of poly-proline tracts. Increased occupancy manifests at CCA and less at CCC. e) Adenosine-ending codons show specific translation defects induced by the combined ProArg-free DFMO treatment when the transcriptome-wide relative ribosome occupancy is compared to control diet. f) Schematic showing two mechanisms of polyamine depletion therapy. Only combined treatment induces hallmarks of eIF5A hypusination deficiency and boosts previously unknown codon specific translation defects induced by polyamine depletion. As described in figure 3, data in B-F are from TH-MYCN mouse model. In vivo treatment with DFMO is 1% in the drinking water. For all Mean, *n* = 5. Abbreviations: CD, control diet; ProArg-free, proline and arginine depleted diet; DFMO difluoromethylornithine.

In addition to its global functions, hypusinated eIF5A has been implicated in facilitating peptide bond formation involving repetitive instances of the amino acid proline, termed poly-proline tracts. Due to its reactive amine localized within a ring structure, proline is a poor peptidyl acceptor^32^. We thus evaluated relative ribosome occupancy at poly-proline tracts, with high occupancy indicating slow decoding by ribosomes, either due to uncharged proline tRNA or eIF5A deficiency. Increased occupancy was observed at poly-proline tracts within ProArg-free DFMO-treated tumors (Fig.4c), but not at proline codons upon diet treatment alone (Extended Data Fig.7d). Collectively, these data argue for the combined diet-drug therapy impairing translation by depleting spermidine to levels low enough to impair eIF5A hypusination.

## Polyamine depletion causes defective decoding of adenosine-ending codons

Further analysis of the ribosome profiling data revealed a surprising aspect of the diet-drug combo-induced stalling at poly-proline tracts: ribosome stalling was observed predominantly at one of the four proline codons (CCA). There was less for CCC and no pausing for CCG or CCT. This indicated an additional unanticipated level of translation regulation at the codon, rather than the amino acid level. Broadening the analysis to all proline codons (i.e., including those not in poly-proline tracts) showed the same phenomenon: selected stalling at CCA codons (Fig.4d).

Therefore, we next assessed the global effect of combined diet-drug treatment across individual codons at high resolution. Strikingly, ribosome pausing depended strongly on the codon type (as opposed to amino acid identity). When sorting individual codons by relative occupancy, a primary factor determining translation speed was the nucleotide at the third position. Whereas adenosine-ending codons showed increased occupancy (i.e., stalling) in ProArg-free DFMO tumors, occupancy at guanosine-ending codons was markedly decreased (Fig.4e and Extended Data Fig.7e). The same phenotype, with a reduced effect size, was shown for the DFMO monotherapy group, suggesting a previously unknown role of polyamines (rather than low arginine or proline) in the decoding of codons with adenosine at the ending position, also called the wobble position (Fig.4f and Extended Data Fig.7f). While interactions of polyamines with ribosomes and tRNAs have been reported^33,34^ and documented to stimulate translation *in vitro*^35^, the codon resolution phenotype has not previously been described (Fig.4f).

## Translational pausing at adenosine-ending codons rewires the proteome

To better understand the phenotypic consequences induced by this specific translation defect, we integrated RNA-seq, Ribo-seq, and proteomics (Fig.5a and Extended Data Fig.8a-d). The main driver of transcriptional changes was DMFO. The ProArg-free diet itself induced only minor such changes, and transcriptional changes strongly overlapped between DFMO alone and the combined diet-drug treated tumors. Ribo-Seq and proteomics, in contrast, revealed distinct changes in tumors under combined ProArg-free DFMO therapy. The effect was independent from proline or poly-proline abundance in the proteins (Extended Data Fig.8), and instead correlated with genes high in adenosine-ending codons.

**Figure 5:**
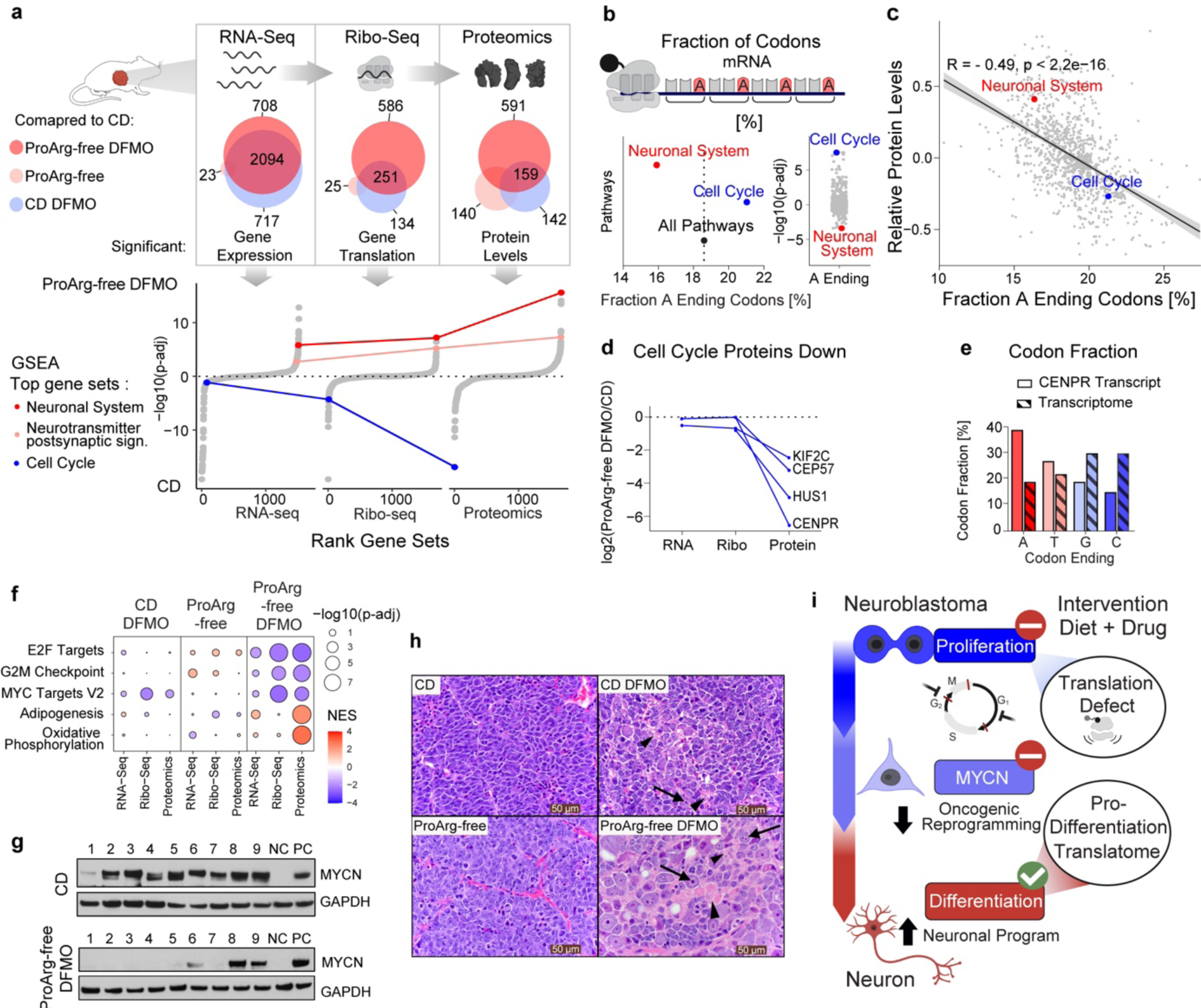
Regulation of translation by polyamine depletion is driven by fractional codon content. a) Venn diagram displaying the number of significant changes per treatment group across the layers of protein biosynthesis: gene expression (RNA-Seq), gene translation (RNA-Seq) and protein levels (Proteomics). Each group was compared to CD by differential analysis. Corresponding enrichment analysis across omics layers comparing ProArg-free DFMO to CD using the Reactome gene sets. All pathways are depicted, ranked by significance and signed. The most down- and upregulated sets are ‘cell cycle’ and ‘neuronal system’ the protein level, respectively, and are connected across the omics layers. b) Average fraction (right) of codons ending with adenosine (A-ending) across all genes or selected pathways or their enrichment based on ranking genes by the fraction of A-ending codons across all Reactome pathways (left). Pathways positively enriched (positive p-adj) have higher A-ending fraction whereas negatively enriched have a lower fraction. c) The percentage of A-ending codons correlates with the average protein levels across Reactome pathways. Pathways with an increasing fraction of A-ending codons have lower protein levels in ProArg-free DFMO compared to CD (log2 fold change). d) Fold change across omics layers of top downregulated cell cycle proteins indicates predominant change on the protein level. e) Percentage of codons with the respective nucleotide at the ending position in the *Itgb3pg* gene (CENPR protein) compared to the transcriptome background. f) Gene set enrichment analysis across omics layers in all three treatment groups using the Hallmark gene set. Only in the ProArg-free DFMO combined treatment group a significant effect is achieved when compared to CD. The major effect is on the translation and protein level. Displayed are the five top enriched sets (complete in Extended Data Fig.10c). Size denotes significance and color normalized enrichment score, with red showing enriched in the intervention group (CD DFMO, ProArg-free or ProArg-free DFMO) and blue in CD. g) Western blot analysis of MYCN in tumors from CD and ProArg-free DFMO arms. NC negative control: CHLA20 neuroblastoma cell line (non-amplified) and PC positive control: IMR5 neuroblastoma cell line (*MYCN* amplified). GAPDH loading control. i) Representative H&E-stained sections. CD and ProArg-free show undifferentiated primitive neuroblasts, absent neuropil, and prominent mitotic figures. CD DFMO shows poorly differentiated primitive neuroblasts with scant neuropil (arrowhead) and foci of cytodifferentiation (<5% differentiating, arrow). ProArg-free DFMO tumors show high fractions of differentiating neuroblasts (>5% differentiating) with increased cytoplasmic to nuclear ratio (arrow) and abundant neuropil (arrowhead). j) Summary of treatment effects. While cell cycle and MYCN programs are downregulated on the proteome due to translation inhibition, immature cancer cells are driven into neuronal differentiation. For a,c,d and f: RNA-Seq: ProArg-free DFMO *n* = 5; CD *n* = 4, Ribo-Seq: *n* = 5, Proteomics: ProArg-free DFMO *n* = 6; CD *n* = 5. Abbreviations: CD, control diet; ProArg-free, proline arginine free diet; DFMO, difluoromethylornithine. Scale bar= 50*μ*m.

Notably, upon combined ProArg-free DFMO treatment, gene sets linked to neuronal differentiation and neuronal cell identity were the most upregulated on the protein level. In contrast, ‘cell cycle’ was the most downregulated, despite having unchanged RNA expression (Fig.5a lower part and Extended Data Fig.9a). We next assessed whether the slow translation of adenosine-ending codons relates to proteome reprogramming. Indeed the ‘cell cycle’ gene set contained a higher frequency of adenosine-ending codons affected by ribosome pausing, as compared to the exome average. Exome-wide, pathways associated with cell cycle programs were the most enriched in adenosine-ending codons^36^. Neuronal differentiation associated gene sets were at the opposite end of the spectrum (Fig.5b and Extended Data Fig.9b-c). The higher the fraction of adenosine-ending codons in a pathway, the lower their respective protein fold change or gene set enrichment (Fig.5c and Extended Data Fig.10d).

Beyond the general reduction of cell cycle proteins, four mitosis related proteins stood out with the strongest downregulation under combined diet-drug treatment (Extended Data Fig.9e-f). All four showed at least several-fold downregulation on the protein level without changes in gene expression (Fig.5d). *Itgb3pg,* the gene encoding for the strongest affected protein CENPR (centromere protein R), showed a remarkably shifted distribution in codon preference with 38.6 % ending on adenosine, as opposed to 18.8% across all protein coding transcripts (Fig.5e and Extended Data Fig.9g-h). The ribosome distribution along the CENPR coding sequence confirmed preferential pausing at adenosine-ending codons upon ProArg-free DFMO treatment as indicated by an increased ribosomal pausing sum at adenosine-ending codons and reduced pausing at guanosine-ending ones (Extended Data Fig.9i-j). Similar pausing was observed for CEP57 and KIF2C (Extended Data Fig.9k-l). Thus, polyamine depletion by a diet-drug combination induces ribosome stalling at adenosine-ending codons. Due to such codons being selectively enriched in cell-cycle genes and depleted in differentiation genes, this reprograms the proteome in a manner that favors differentiation.

## Polyamine depletion-mediated proteome rewiring induces neuroblastoma differentiation

We hypothesized that the shift away from cell-cycle genes and towards neuronal differentiation genes would reprogram neuroblastoma tumors in a manner that slows proliferation and induces differentiation. Ki67 staining confirmed a decreased number in actively cycling cells under ProArg-free DFMO treatment (Extended Data Fig.10a). On western blot, proliferating cell nuclear antigen (PCNA, a proliferation marker) showed decreased levels selectively in the ProArg-free DFMO treatment group (Extended Data Fig.10b). Furthermore, cell growth activation signatures including MYC and E2F targets were impaired (Fig.5f and Extended Data Fig.10c).

Given the essential role *of MYCN* in the development and maintenance of neuroblastoma, we then asked whether the loss of ‘MYC targets’ indicates a disruption of the core oncogenic regulatory circuit^37–39^ (Fig.10f). Both *MYCN* mRNA expression and MYCN protein were preferentially downregulated in tumors under combined ProArg-free DFMO treatment (Fig.5g and Extended Data Fig.10d-e). Other core transcription factors were also suppressed supporting a broad disruption of the MYCN-driven core regulatory circuit (Extended Data Fig.10e). Evaluating tumor differentiation status according to clinical pathological criteria showed tumors in the CD and ProArg-free groups as uniformly undifferentiated (<5% cytologic differentiated without neuropil). In contrast, upon polyamine depleting treatment a strong differentiation phenotype was observed with one third of CD DFMO treated tumors being differentiated (>5%), and two thirds of ProArg DFMO treated tumors differentiating or partially differentiated with abundant neuropil (a feature of neural differentiation; Fig.5h and Extended Data Fig.10f). Thus, combining a proline/arginine-free diet with DMFO therapy led to marked reductions in polyamines, inducing selective translation defects that suppressed tumor cell proliferation and induced tumor differentiation (Fig.5i).

## Discussion

We find that neuroblastoma, a highly malignant childhood cancer, is vulnerable to polyamine depletion mediated by the combination of a diet free of proline and arginine to deplete the polyamine precursor ornithine and pharmacological inhibition of the committed step of polyamine synthesis, ornithine decarboxylase, with DFMO. The diet-drug combination enhanced polyamine depletion and exerted strong anti-cancer effect in a highly lethal murine neuroblastoma model, with apparent cures in some mice.

Use of defined diet-drug combinations is emerging as a clinically viable strategy for cancer treatment^41,42^. Ketogenic diet can synergize with both classical chemotherapy and targeted agents, and can be achieved, with proper support, by patients making careful food choices^43–45^ (NCT05300048, NCT01535911). Diets lacking certain amino acids require laboratory formulation, but show acceptable taste and desired metabolic effects in humans and are also entering cancer efficacy trials^46,47^ (NCT05078775). Such diets have related effects to enzyme-based treatments that catabolize particular amino acids, such as asparaginase, a long-standing standard of care for childhood leukemias^48^. In some cases, however, the diets have important advantages. For example, dietary arginine depletion decreases both arginine and ornithine, while arginase therapy^49^ (which is in clinical trials but has yet to show benefits in humans) depletes arginine by converting it into ornithine, which accordingly goes up. Thus, only the dietary approach is suitable for achieving ornithine depletion and effective combination with DMFO to deplete polyamines.

Arrested cellular differentiation by retained embryonal gene expression circuits is a hallmark of pediatric cancers^50,51^. Neuroblastoma is a prime example with hyperactive MYC(N) driving embryonal programs^37,39,40,52^. Differentiation induction through transcriptional reprogramming is a validated therapeutic strategy, exemplified by the nuclear hormone receptor agonist retinoic acid^53,54^. We provide evidence for the feasibility of triggering differentiation in pediatric cancers also through proteome reprogramming. Specifically, we find that polyamine depletion promotes translation of pro-differentiation proteins and suppresses that of cell-cycle ones, leading to neuronal differentiation of neuroblastoma. Whether similar benefits could be achieved in other MYC-driven cancers merits investigation.

The selective effect of polyamine depletion on translation of certain gene sets was unexpected. The underlying biochemistry involves polyamine deficiency impairing translation of codons with adenosine at the wobble base, and such codons being preferentially enriched (or depleted) in different gene sets. This biochemistry raises the possibility that codon usage has evolved in tandem with metabolism, such that metabolic limitation acts on translation of specific codons to rewire the proteome. Specifically, our data point to polyamine depletion working through adenosine-ending codons to suppress proliferation and promoting differentiation, a program that evolved, perhaps to support proper developmental decisions, and can be hijacked by the combination of diet and pharmacology to treat neuroblastoma.

## Acknowledgements

We are grateful to members of the Rabinowitz, Hogarty and Morscher groups for scientific discussions and insights. We thank Gernot Poschet and Glynis F. Klinke at the Metabolomics Core Technology Platform of the Excellence Cluster “Cell Networks” (University of Heidelberg; ZUK49/2) for support with metabolite analyses, Ngo Quy Ai for her support with data analysis, Andrea Franson and Ocean Malka for technical assistance. We thank Alexander Kowar from the Translational Control and Metabolism group at the DKFZ for methodological discussions. We thank and acknowledge the staff at the Princeton and CHOP animal facility core. Mass spectrometry analyses were performed by the Proteomics Research Infrastructure (PRI) at the University of Copenhagen (UCPH), supported by the Novo Nordisk Foundation (NNF) (grant agreement number NNF19SA0059305). We thank Rita Colaco for support with proteomic data analysis, Paola Pisano, and Emma Ahrman for proteome analyses, and Katarzyna Wozniak for technical assistance. Cell lines were provided by the Children’s Oncology Group Cell Culture Repository (cccells.org) and primary neuroblastoma samples by the Children’s Oncology Group Biospecimen Repository. M.D.H. was supported by a US Department of Defense Team Translational Science Award (CA170257). This work was supported by the US National Institutes of Health (NIH Pioneer award DP1DK113643 to J.D.R and R01CA163591 to J.D.R. and E.W.). S.C. and R.J.M. were supported by grants from the NOMIS Foundation, Holcim-Stiftung Wissen, Gertrud-Hagmann-Stiftung für Malignom-Forschung, EMDO-Stiftung and Heidi Ras Grant of the FZK University Children’s Hospital Zürich.

## Author contributions

Conceptualization and methodology, S.C., E.W., J.D.R., M.D.H. and R.J.M.; investigation and data analysis, S.C., L.Y., M.M., C.S.T., W.L., C.E., G.E.A., O.O.P, L.Z., A.V., Y.L., O.H.G., L.S., M.W., M.D.H., R.J.M.; writing – original draft, S.C. and R.J.M; writing – review & editing, S.C., E.W., J.D.R., M.D.H. and R.J.M.; resources, E.W., J.D.R., M.D.H. and R.J.M.; supervision and funding acquisition, E.W., J.D.R., M.D.H. and R.J.M.;

## Methods

### Primary neuroblastoma patient samples

Flash-frozen primary neuroblastoma tumor samples were provided by the Children’s Oncology Group (COG) under study number ANBL16B2-Q. International Neuroblastoma Pathology Classification (INPC) histologic parameters. Histology of poorly differentiated neuroblastoma, *MYCN* amplification status, age, and stage and pathologic classification for every patient was obtained centrally via COG testing and review. Tumor cell content of samples was confirmed as > 80% percent. Patient and tumor characteristics are given in Table S1. Water soluble metabolites were extracted and analyzed as described below.

### Mouse models

Animal studies followed protocols approved by the Princeton University and Children’s Hospital of Philadelphia Institutional Animal Care and Use Committees. For xenografts, cancer cell lines were grown in RPMI supplemented with 10% FBS and 0.01 % insulin/transferrin solution. Cell lines were provided by the Children’s Oncology Group Cell Culture Repository: LA-N-5, SMS-SAN, CHLA-90 and SK-N-SH. All cell lines repeatedly tested negative for Mycoplasma. Subcutaneous xenografts were established on 6-week old female CD1-nu mice by injection of 100ul 50/50 RPMI/Matrigel solution containing 10^6 cells of the respective cell line. The TH-*MYCN* mouse model was used as a primary model to investigate the functional changes of metabolism driven by MYCN. 129×1/SvJ mice transgenic for the TH-*MYCN* construct^10^ were originally obtained from Bill Weiss (University of California, San Francisco). TH-*MYCN* hemizygous mice were bred and litters randomized to assigned therapy. Mice were genotyped from tail-snip-isolated DNA using qPCR^23^ and only transgene homozygous mice (TH-*MYCN*+/+) were included in these studies. In this model, MYCN expression is targeted to the murine neural crest under the tyrosine hydroxylase promoter recapitulating the lethal hallmark features of human neuroblastoma.

### Metabolite extraction from tissue, tumors and serum

Tissues and tumors were collected from mice in fed state and immediately clamped into liquid nitrogen using Wollenberger clamp. All tissues were stored in −80°C. Frozen tissues were transferred into 2 ml Eppendorf tubes, which were precooled on dry ice, and pulverized using Cyromill. The resulting tissue powder was weighed (around 10mg) and mixed well by vortexing in extraction buffer (40 µL extraction buffer per mg tissue). The extraction solution was neutralized with NH_4_HCO_3_ as above and centrifuged in a microfuge at maximum speed for 30 min at 4°C. Supernatant was transferred to LC-MS vials for analysis. Blood samples were drawn from mouse tail veins using a microvette and kept on ice. After centrifugation (10 min, benchtop microfuge maximum speed, 4°C), serum was collected in a 1.5ml tube and store in −80 °C. 5 µl of serum were mixed with 200 µl of extraction buffer (40:40:20 acetonitrile: methanol: water with 0.5% formic acid) and neutralized with 15% NH_4_HCO_3_. After centrifugation (30 min, benchtop microfuge maximum speed, 4°C), supernatant was transferred into LC-MS vial for analysis.

### Metabolite measurement by liquid chromatography-mass spectrometry

Metabolomics was performed on the following systems. A quadrupole-orbitrap mass spectrometer (Q Exactive, Thermo Fisher Scientific), operating in positive or negative mode was coupled to hydrophilic interaction chromatography (HILIC) via electrospray ionization^16^. Scans were performed from m/z 70 to 1000 at 1 Hz and 140 000 resolution. LC separation was on a XBridge BEH Amide column using a gradient of solvent A (20 mM ammonium acetate, 20 mM ammounium hydroxide in 95:5 water:acetonitrile, pH 9.45) and solvent B (acetonitrile). Flow rate was 150 mL/min. The LC gradient was: 0 min, 85% B; 2 min, 85% B; 3 min, 80% B; 5 min, 80% B; 6 min, 75% B; 7 min, 75% B; 8 min, 70% B; 9 min, 70% B; 10 min, 50% B; 12 min, 50% B; 13 min, 25% B; 16 min, 25% B; 18 min, 0% B; 23 min, 0% B; 24 min, 85% B. Autosampler temperature was 5°C, and injection volume was 5-10 uL. Complementary, primary samples analyzed on an Exactive (Thermo Fisher Scientifc) operating in negative ion mode^55^. Liquid chromatography separation was achieved on a Synergy Hydro-RP column (100 mm × 2 mm, 2.5 µm particle size, Phenomenex, Torrance, CA), using reversed-phase chromatography with the ion pairing agent tributylamine in the aqueous mobile phase to enhance retention and separation. An adaptive scan range was used with an m/z from 85-1000. Resolution was 100 000 at 1 Hz. The total run time is 25 min with a flow rate at 200 µL/min. Solvent A is 97:3 water/methanol with 10 mM tributylamine and 15 mM acetic acid; solvent B is methanol. The gradient is 0 min, 0% B; 2.5 min, 0% B; 5 min, 20% B; 7.5 min, 20% B; 13 min, 55% B; 15.5 min, 95% B; 18.5 min, 95% B; 19 min, 0% B; 25 min, 0% B.

### Mass spectrometry analysis

Metabolomics data analysis was performed using ElMaven software (https://github.com/ElucidataInc/ElMaven). For labelling experiments, correction for natural abundance of 13C was performed using Accucor (https://github.com/XiaoyangSu/AccuCor).

### Infusion studies and isotope tracing in TH-*MYCN* mice

TH-*MYCN* mice were housed in groups and food was supplied without restriction to guarantee sufficient supply. Mice weights were recorded every day. During experiments mice were freely moving and tissues and serum was analyzed following the above mentioned method. Tumor and inter organ cooperativity in proline, arginine and ornithine biosynthesis was analyzed on whole body level. The mice were on normal light cycle (6 AM – 6 PM). In vivo infusion was performed on 6-7-week old normal TH-*MYCN* mice pre-catheterized on the right jugular vein and ^13^C metabolite tracers were infused for 2.5-5 h to achieve isotopic pseudo-steady state. The mouse infusion setup included a tether and swivel system, connecting to the button pre-implanted under the back skin of mice. Mice were fasted from 9:00 am to 2 pm and infused from 2pm till 4:30 pm. Tracers were dissolved in saline and infused via the catheter at a constant rate (0.1 µl/min/g mouse weight) using a Just infusion Syringe Pump. 100mM [U-^13^C]glutamine was dissolved and infused for 2.5 hours, 40mM [U-^13^C]arginine for 5 hours, 200mM [U-^13^C]glucose for 5 hours, 10mM [U-^13^C]proline for 5 hours and 5mM [U-^13^C]ornithine for 5 hours. At the end of infusion, mice were dissected and tissues were clamped in aluminum foil and stored in liquid nitrogen.

### Intervention study in TH-*MYCN* mice

Arginine and Proline free diet was purchased from TestDiet® Baker under the catalog number 1812426 (5CC7) for control diet and 1816284-203 (5WYF) for arginine / proline free diet. Detailed makeup is given in Table S2. The ODC-inhibitor, DFMO, was obtained from Pat Woster (Medical University of South Carolina). DFMO was dissolved in drinking water and supplied to mice *ad libitum* at a dose of 1% in the drinking water. *Survival end-point*: TH-*MYCN*+/+ mice were randomized to DFMO or no DFMO at birth, and diets changed to amino acid-based control diet or arginine / proline free diet at day 21, per treatment assignment. Mice were weighed and assessed for tumor growth and symptoms, at least thrice weekly. Mice were euthanized for pre-defined humane endpoints related to overall health or tumor burden. For metabolomics studies TH-*MYCN*+/+ mice were treated as above, serum was obtained at day 43 (+/− 2 days) and tumors and organs harvested at that time if tumor present, or delayed to the earliest time a tumor became palpable. Time to tumor harvest depended on the treatment group (Extended Data). For homogenous timing of tumor harvest TH-*MYCN*+/+ mice were assigned to delayed DFMO and diet change at day 28 to enable uniform assessment of metabolomic changes in tumors (Main Figure).

### Polyamine quantification

Polyamine concentrations were quantified using the AccQ-Tag fluorescence dye (Waters) as described by Yang et al.^56^. Derivatives were separated on an Acquity BEH C18 column (150 mm x 2.1 mm, 1.7 µm, Waters) by reverse phase UPLC (Acquity H-class UPLC system, Acquity FLR detector, Waters). The column was equilibrated with buffer A (140 mM sodium acetate pH 6.3, 7 mM triethanolamine) at a flow rate of 0.45 ml min-1 and heated at 42°C. Pure acetonitrile served as buffer B. The gradient was produced by the following concentration changes: 1 min 8% B, 7 min 9% B, 7.3 min 15% B, 12.2 min 18% B, 13.1 min 41% B, 15.1 min 80% B, hold for 2.2 min, and return to 8% B in 1.7 min. Chromatograms were recorded and processed with the Empower3 software (Waters). For acetylated polyamines a MS/MS method was used. In brief, a Waters Acquity I-class Plus UPLC system (Binary Solvent Manager, thermostatic Column Manager and FTN Sample Manager) (Waters, USA) coupled to an QTRAP 6500+ (Sciex, USA) mass spectrometer with electrospray ionization (ESI) source was used. Data acquisition was performed with Analyst (Sciex, USA) while data quantification was performed with the SciexOS software suite (Sciex, USA). Chromatography was made on an Acquity HSS T3 column (150 mm x 2.1 mm, 1.7 µm, Waters) kept at 20 °C and a flow rate of 0.3 ml/min. Eluent A consisted of water with 0,1% formic acid and eluent B in ACN with 0,1% formic acid. Gradient elution consisted in changing %B as follows: 0-1 min 0% 5 min 20%; 5,5-7,5 min 100%, and 8-10 min 0%. The ion source settings were as follow: curtain gas: 30 psi; collision gas: low; ion spray: 4500 V; source temperature: 500°C; ion source gas 1: 40 (GS1) and ion source gas 2: 50 (GS2). All compounds were measured in positive electrospray ion mode.

### Metabolomics and MYCN transcriptional activity 180 cancer cell lines

Metabolomics of 180 cancer cell lines has been obtained from Cherkaoui et al.^57^. Association between metabolites levels of core metabolic pathways and MYCN transcriptional activity have been obtained from interactive dashboard (https://cancer-metabolomics.azurewebsites.net/page2).

### Gene expression analysis in neuroblastoma tumors

Gene expression profiles of 649 neuroblastoma tumors^21^ were obtained from R2 (R2: Genomics Analysis and Visualization Platform (http://r2.amc.nl)). Differential expression analysis between MYCN amplification status was performed using the Bioconductor package ‘limma’^58^ (version 3.40.6).

### Gene expression analysis in neuroblastoma cell lines

Expression profiles of 39 neuroblastoma cell lines^59^ were obtained from Gene Expression Omnibus. Differential analysis was performed using the Bioconductor package ‘limma’ (version 3.40.6) and by comparing MYCN amplification status provided in the study.

### RNA and ribosome sequencing

#### Isolation of total RNA, library preparation and sequencing

Total RNA were isolated from the same extracts, that were used to obtain RPF (bellow, Ribo-Seq). 3 volumes of QIAzol® (Qiagen, Cat. No. 79306) were added to 80 µl of cell extracts, mixed thoroughly and proceed to RNA purification with Direct-Zol RNA Mini Prep Plus kit. RNA were sent to Genomic Platform (UNIGE) for stranded mRNA libraries preparation. Libraries were sequenced on an Illumina NovaSeq 6000, SR 100 bp, 10 libraries in 1 pool.

### Ribo-Seq

Mouse tumors were mechanically disrupted in liquid nitrogen and homogenized in a lysis buffer (LB, 50 mM Tris, pH 7.4, 100 mM KCl, 1.5 mM MgCl2, 1.0% Triton X-100, 0.5% Na-Deoxycholate, 25 U/ml Turbo DNase I, 1mM DTT, 100 µg/ml cycloheximide, and Protease inhibitors) 3 ml of LB per 1 g of tissue. To obtain ribosome footprints 0.12 ml of total extracts containing 300 µg of total RNA were treated with RNAse I (Epicentre) (25U/1 mg of total RNA), for 45 min at 20°C with slow agitation. 10 ml SUPERaseIn RNase inhibitor was added to stop nuclease digestion. Monosomes were isolated using S-400 columns. For isolation of mRNA protected fragments (RPF) 3 volumes of QIAzol® were added to the S-400 eluate, mixed thoroughly and proceed to RNA purification with Direct-Zol RNA Mini Prep Plus kit.

RPF libraries were prepared as described^27,60^. Briefly, RPFs (25-34 nt) were size-selected by electrophoresis using a 15% TBE-Urea polyacrylamide gel electrophoresis (PAGE) and two RNA markers, 25-mer (5’ AUGUACACGGAGUCGAGCACCCGCA 3’) and 34-mer (5’AUGUACACGGAGUCGAGCACCCGCAACGCGAAUG 3’). After dephosphorylation with T4 Polynucleotide Kinase (NEB, #M0201S) the adapter Linker-1 (5’ rAppCTGTAGGCACCATCAAT/3ddC/ 3’) was ligated to the 3′ end of the RPF using T4 RNA Ligase 2. Ligated products were purified using 10% TBE-Urea PAGE. Ribosomal RNA was subtracted using RiboCop rRNA Depletion Kit V2 H/M/R. The adapter Linker-1 was used for priming reverse transcription (RT) with the RT primer Ni-Ni-9 (5’AGATCGGAAGAGCGTCGTGTAGGGAAAGAGTGTAGATCTCGGTGGTCGC_5_CACTCA_5_TTCAGACGTGTGCTCTTCCGATCTATTGATGGTGCCTACAG 3’) using ProtoScript® II Reverse Transcriptase. RT products were purified using 10% TBE-Urea PAGE. The cDNA was circularized with CircLigase™ II ssDNA Ligase. The final libraries were generated by PCR using forward index primer NI-N-2 (5’ AATGATACGGCGACCACCGAGATCTACAC 3’) and one of the reverse index primers. Amplified libraries were purified using 8% TBE-PAGE and analyzed by Qubit and TapeStation. Libraries were sequenced on an Illumina NovaSeq 6000, SR 100 bp, 4 libraries in 1 pool.

### RNA-Seq mapping

Fastq files were adaptor stripped using cutadapt with a minimum length of 15 and a quality cutoff of 2 (parameters: -a CTGTAGGCACCATCAAT –minimum-length = 15 –quality-cutoff = 2). Resulting reads were mapped, using default parameters, with HISAT2^61^, using a GRCm38, release 101 genome and index. Differential expression analysis was performed using DESeq2^62^, using a GRCm38, release 101 genome and index.

### Ribo-Seq mapping

Fastq files were adaptor stripped using cutadapt. Only trimmed reads were retained, with a minimum length of 15 and a quality cutoff of 2 (parameters: -a CTGTAGGCACCATCAAT – trimmed-only –minimum-length = 15 –quality-cutoff = 2). Histograms were produced of ribosome footprint lengths and reads were retained if the trimmed size was 28 or 29 nucleotides. Resulting reads were mapped, using default parameters, with HISAT2^61^ using a GRCm38, release 101 genome and index and were removed if they mapped to rRNA or tRNA according to GRCm38 RepeatMasker definitions from UCSC. A full set of transcripts and CDS sequences for Ensembl release 101 was then established. Only canonical transcripts [defined by knownCanonical table, downloaded from UCSC] were retained with their corresponding CDS. Reads were then mapped to the canonical transcriptome with bowtie2 using default parameters.

### Ribo-Seq analysis

The P-site position of each read was predicted by riboWaltz^63^ and confirmed by inspection. Counts were made by aggregating P-sites overlapping with the CDS and P-sites Per Kilobase Million (PPKMs) were then generated through normalizing by CDS length and total counts for the sample. Differential expression and translational efficiency analysis was performed using DESeq2^62^. All metagenes, stalling and Ribosome Dwelling Occupancy (RDO) analyses are carried out on a subset of expressed canonical transcripts which had PPKM values greater than 1 across all samples (10366 total). Within these, P-site depths per nucleotide were normalized to the mean value in their respective CDS. For metagenes around codon types, the mean of these normalized values is taken for each codon within 90 nucleotides of every instance of that codon. For RDO calculation for a given type of codon, the mean of these normalized values is taken over all instances of that codon, then these are compared using a log2FC-ratio between conditions. To assess relative pausing, P-site depths normalized to the CDS mean were compared at each codon position in the CDS. A value of 1 was added to these normalized depths and a log2FC ratio was taken pairwise between conditions. To compare effects of different codon ending bases, the resulting values were separated by the ending base of each codon and plotted across their respective positions in the CDS. The relative pausing sum for each ending A, T, G or C is then the sum of these values for every codon containing the respective ending codon across the CDS. The fraction of nucleotides at ending codons were evaluated from extracting the codons for the CDS of each gene using GRCm38, release 101. Pathway level fractions were computed using the average of each gene contain in the pathway.

### Immunoblots

Harvested TH-*MYCN* tumors were clamped and flash-frozen in liquid nitrogen. After this mechanical dissociation, crude protein extraction was obtained by lysis with CHAPS buffer (10mM HEPES, 150mM NaCl, 2% CHAPS) with fresh protease inhibitor and phosphatase inhibitor. This protein lysate (25 micrograms) was electrophoresed through a 5-10% Tris-Glycine gel and immunoblotted using antibodies to MYCN (1:200, Santa Cruz) and PCNA (1:1000, Cell Signaling Technologies).

### Isoelectric Focusing Blots

Crude protein extraction obtained as described above was electrophoresed through a slab isoelectric focusing gel (pH 3-7, Invitrogen Novex EC66452) employing freshly made cathode and anode buffers (Novex). The gel was transferred to a PVDF membrane and transferred using the iBlot transfer unit prior to blocking in buffer according to manufacturer’s recommendations for iBind. The iBind was then assembled with a probe against eIF5A (BD Laboratories, 1:3000) and incubated for at least 2.5 hours prior to development.

### Histology

Harvested TH-*MYCN* tumors were preserved in 10% formalin and embedded in paraffin blocks. Slides were cut and then stained with hematoxylin and eosin (H&E). These slides were reviewed by a pathologist blinded to the treatment groups, and tumors were scored according to 1) differentiation status, 2) neuropil presence or absence and relative abundance, and 3) evidence of global or localized necrosis. Slides were then scanned and re-reviewed by the same pathologist.

### Immunohistochemistry

Slides of formalin fixed, paraffin embedded tumors (prepared as above) were stained with *MYCN* (1:100, Abcam) and Ki67 (1:200, Abcam). Staining was performed on a Bond Max automated staining system (Leica Microsystems). The Bond Refine staining kit (Leica Microsystems DS9800) was used. The standard protocol was followed with the exception of the primary antibody incubation which was extended to 1 hour at room temperature. Antigen retrieval was performed with E2 (Leica Microsystems) retrieval solution for 20min. After staining, slides were rinsed, dehydrated through a series of ascending concentrations of ethanol and xylene, then cover-slipped. Stained slides were then digitally scanned at 20x magnification on an Aperio CS-O slide scanner (Leica Biosystems). These slides were reviewed by a pathologist blinded to the treatment groups and scored according to differential MYCN expression and Ki67 staining, respectively.

### Proteomics

#### Total proteome sample preparation

Tissues were disrupted by grinding in frozen state and lysed in lysis buffer (20 mM HEPES pH 7.2, 2% SDS). Proteins extracts were diluted 1:1 with 2x SDS buffer (10% SDS, 100 mM Tris pH 8.5), boiled for 10 min at 95 °C, reduced with 5 mM (final) Tcep for 15 min at 55 °C, and alkylated with 20 mM (final) CAA for 30 min at room temperature. Proteins were acidified by addition of 3% (final) perchloric acid, followed by addition of seven volumes of Binding buffer (90% MeOH, 100 mM TEAB). Samples were loaded on S-trap columns and processed on a Resolvex A-200 positive pressure unit (Tecan). Samples were washed 1x with Binding buffer, 3x with 50% MeOH/50% CHCl3 and 2x with Binding buffer. 150 µL of digestion buffer (TEAB 50 mM) containing Trypsin 1:10 (wt:wt, enzyme:protein) and Lys-C mix 1:50 (wt:wt, enzyme:protein) was added and incubated for 1h at 37C. 100 µL of digest buffer was added, and incubated over-night. Peptides were eluted with 80 µL 0.2% aqueous formic acid followed by 80 µL of 50% ACN containing 0.2% formic acid. Peptides were diluted 1:1 with STOP buffer (PreOmics) and purified over iST positive pressure plates (PreOmics) according to the manufacturer’s instructions.

### Nanoflow LC-MS/MS measurements for proteomes

Peptides were separated on an Aurora (Gen3) 25 cm, 75 µM ID column packed with C18 beads (1.7 µm) (IonOpticks) using a Vanquish Neo (Thermo Fisher Scientific) UHPLC. Peptide separation was performed using a 90-minute gradient of 2-17% solvent B (0.1% formic acid in acetonitrile) for 56 min, 17-25% solvent B for 21 min, 25-35% solvent B for 13 min, using a constant flow rate of 400 nL/min. Column temperature was controlled at 50°C. MS data were acquired with a timsTOF HT (Bruker Daltonics) in diaPASEF mode. MS data were collected over a 100-1700 m/z range. During each MS/MS data collection each PASEF cycle was 1.8 seconds. Ion mobility was calibrated using three Agilent ESI-L Tuning Mix ions 622.0289, 922.0097 and 1221.9906. For diaPASEF we used the long-gradient method which included 16 diaPASEF scans with two 25 Da windows per ramp, mass range 400.0-1201.0 Da and mobility range 1.43-0.6 1/K0. The collision energy was decreased linearly from 59 eV at 1/K0 = 1.6 to 20 eV at 1/K0 = 0.6 Vs cm-2. Both accumulation time and PASEF ramp time was set to 100 ms.

### MS data quantification

Raw mass spectrometry data were analyzed with Spectronaut (v. 17.1) in directDIA mode with standard settings. Database search included the mouse Uniprot FASTA database.

### Proteomics analyses

Protein intensity values were normalized by log2 transformation and proteins with less than 70% of valid values in at least one group were filtered out. The remaining missing values were imputed using the mixed imputation approach^64^. Briefly, missing values in samples belonging to the same group were imputed with k-nearest neighbors (kNN) if there is at least 60% of valid values in that group, for that protein. The remaining missing values are imputed with the MinProb method (random draws from a Gaussian distribution; width = 0.2 and downshift = 1.8).

### Circulating Turnover Flux (*F*_*circ*_ and 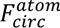)

To measure the circulating turnover flux of a metabolite, we infused [U-^13^C]labeled form of the respective metabolite via the jugular venous catheter. At pseudo-steady state, the fraction of the labeled of mass isotopomer form [*M* + i] of the nutrient in serum is measured as *L*_[*M* + *i*]_, such that i is from 0 to C and C is the total number of carbons in the metabolite. The circulatory turnover flux **F*_circ_* is defined as previously^22^:

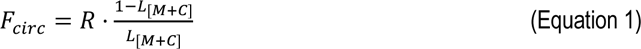

where R is the infusion rate of the labeled tracer. Since the turnover flux is a pseudo-steady state measurement, for minimally perturbative tracer infusions, production flux is approximately equal to consumption flux of the metabolite and thus **F*_circ_* reflects both the circulating production and consumption fluxes of the infused metabolite, in unit of nmol C/min/g. The carbon-atom circulatory turnover flux 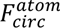 of the nutrient is calculated using:

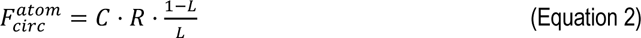

where L is the fraction of labeled carbon atoms in the nutrient:

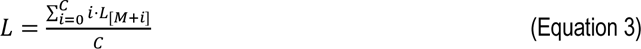

*F_circ_* measures the turnover of the whole carbon skeleton of the molecule, whereas 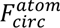 measures the turnover of the carbon atoms in the molecule.

**Normalized metabolite labeling**: When a [U-^13^C]labeled tracer X is infused, the normalized labeling of downstream metabolite Y is defined as 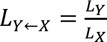, where L_X_ and L_Y_ are the fraction of labeled carbon atoms for metabolite X and Y defined in Equation 3.

### Fractional Contribution to Tissue Metabolites

Direct fractional contribution of each metabolite to other metabolites in a tissue is calculated by setting up the follow set of linear equations:

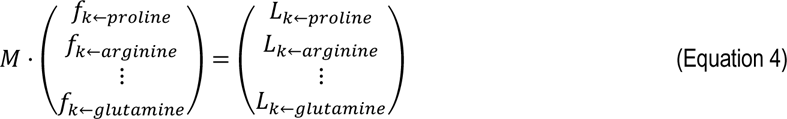

Where *f_k_*_←*i*_ is the fraction of k derived directly from i, M the circulating metabolite interconversion matrix and is taken such that entry (X, Y) represents *L_y_*_←*x*_. Direct contributions to tissue were then calculated by performing an optimization procedure conditional on non-negative values by finding min|| M, f – L || with respect to f such that f > 0. Standard error was estimated using a bootstrapping method (n = 100 simulations) by selecting values for M and L from normal distributions with means and standard deviations equal to calculated values for those parameters based on measured data.

### Circulating Metabolites Flux

The procedure for calculating fluxes between circulating nutrients has been previously described thoroughly in Hui et al.^22^. The input data for this calculation are the inter-labeling matrix M that reflects the extent to which infusion of any nutrient i (of n total nutrients of interest) labels every other circulating nutrient j and the carbon-atom circulatory turnover flux for each circulating nutrient 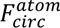. The procedure first uses M to calculate the direct contributions to each nutrient i from all other circulating nutrients j, creating a new n x n matrix N whose entries N_ij_ reflect the direct contributions of circulating nutrient j to circulating nutrient i. It then utilizes the matrix N and 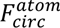 to calculate the direct contributing fluxes from any circulating nutrient to any other circulating nutrient, resulting in a complete determination of the inter-converting fluxes in unit of nmol C/min/g between circulating nutrients. Fluxes were computed using Matlab (v.R2019a).

### Multi-omics Gene Set Enrichment Analysis

All omics were analyzed and visualized in R (Statistical Computing, v4.1.0). Gene set enrichment was performed using fgsea (https://bioconductor.org/packages/release/bioc/html/fgsea.html) using as input gene or protein list rank by relative changes, log2 fold-change of comparison. Gene sets were taken from the Mouse MSigDB Collections, using the gene sets MH: Hallmark and the M2: canonical pathways using the Reactome subset^65^. We used 1,000 permutations of the gene-level values to calculate normalized enrichment scores and statistical significance.

### Statistical information

The number of human and mice in every experiment is recorded in each figure legend. For human tumor measurements, n represents de number of patients. For mice experiments, n represents the number of animals. To determine significance, P-values were computed using an unpaired two-sided Welch’s t-test using the Welch–Satterthwaite equation (not assuming equal variances) unless specified otherwise. Regression between adenosine-ending codon and protein levels were calculated R function ‘stat_cor’ (package ggpubr) to compute Pearson’s r and ‘geom_smooth’ (package ggplot2) using as ‘linear model’ to display the regression line. Statistics were performed using R (Statistical Computing, 4.1.0).

**For metabolomics:** A two-tailed unpaired Welch’s t-test was used to calculate p-values. Metabolomics data were corrected for multiple comparisons using Benjamini–Hochberg’s method, with an FDR cut-off of 0.05 used to determine statistical significance.

**For transcriptomics:** P-values were corrected for multiple hypothesis testing using Benjamini-Hochberg’s method, with an FDR cut-off of 0.05 used to determine statistical significance.

**For proteomics:** Both two-sided Student t-tests were calculated with a permutation-based FDR cut-off of 0.05 and an s0 = 1 if not otherwise declared.

**For mouse survival analyses:** Comparisons of outcome between groups were performed by a two-sided log-rank test, tumor related death is counted as an event, with mice censored at the time of death without tumor on necropsy. Survival was statistically assessed according to the method of Kaplan and Meier^66^ with SEs according to Peto and Peto^67^.

### Data and code availability

The RNA-Seq and Ribo-Seq data are accessible under the accession number GEO: GSE244378

The mass spectrometry proteomics data will be deposited to the ProteomeXchange Consortium via the PRIDE partner repository with the dataset identifier: PXD047396.

Any additional information required to reanalyze the data reported in this manuscript is available from the corresponding author upon request.

## Supplementary Figures

**Extended Data Fig. 1:**
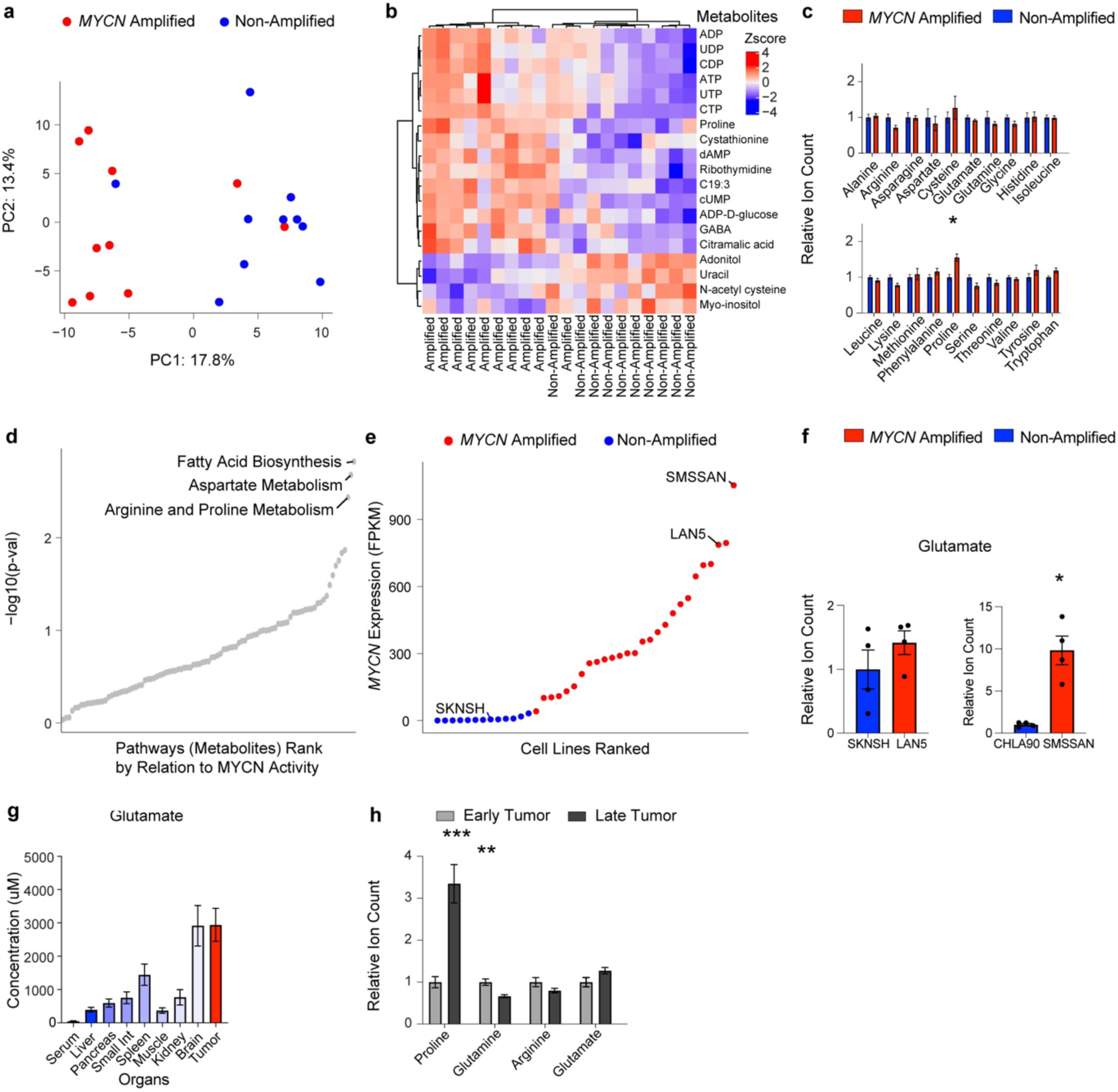
Metabolomic profiling of *MYCN* amplified primary patient tumors and xenografts reveals reprogramming of the arginine-proline-glutamine axis. a) Global metabolomic signatures of primary human neuroblastoma tumors analyzed by principal component analysis, with *MYCN* amplified (red) and non-amplified (blue). b) Heatmap of significantly changed metabolites in primary neuroblastoma tumor samples (q < 0.05), comparing *MYCN* amplified to non-amplified tumors. Unsupervised clustering performed using Ward’s method. c) Levels of all proteinogenic amino acids in primary neuroblastoma tumor tissue. Data in a-c as in Fig.1b with n = 10 each group and corrected p-values, *q < 0.05. Mean ± s.e.m.. d) MYCN transcriptional signatures from gene expression associate with metabolite levels of the arginine and proline metabolic pathway. Data from metabolome profiling of 180 cancer cell lines^57^. e) *MYCN* expression across neuroblastoma cell lines, colored by *MYCN* amplification status based on Harenza et al. ^59^. Cell lines selected for xenografts are depicted by their name. CHLA90, not in this dataset, is a *MYCN* non-amplified line. f) Glutamate levels in contralateral xenografts derived from neuroblastoma cell lines displayed by *MYCN* amplification status. **P < 0.01, two-tailed paired t-test. Mean ± s.e.m., *n* = 4. g) Glutamate levels across organs in the TH-*MYCN* model. Mean ± s.e.m., tumor glutamate *n* = 29, other organs *n* = 8-29. h) Relative metabolite levels across tumors harvested at early (< 50 mm^3^) compared to the late timepoint in the TH-*MYCN* model. **P < 0.01, ***P < 0.001, Two-tailed t-test. Mean ± s.e.m., early *n* = 10, late *n* = 14.

**Extended Data Fig. 2:**
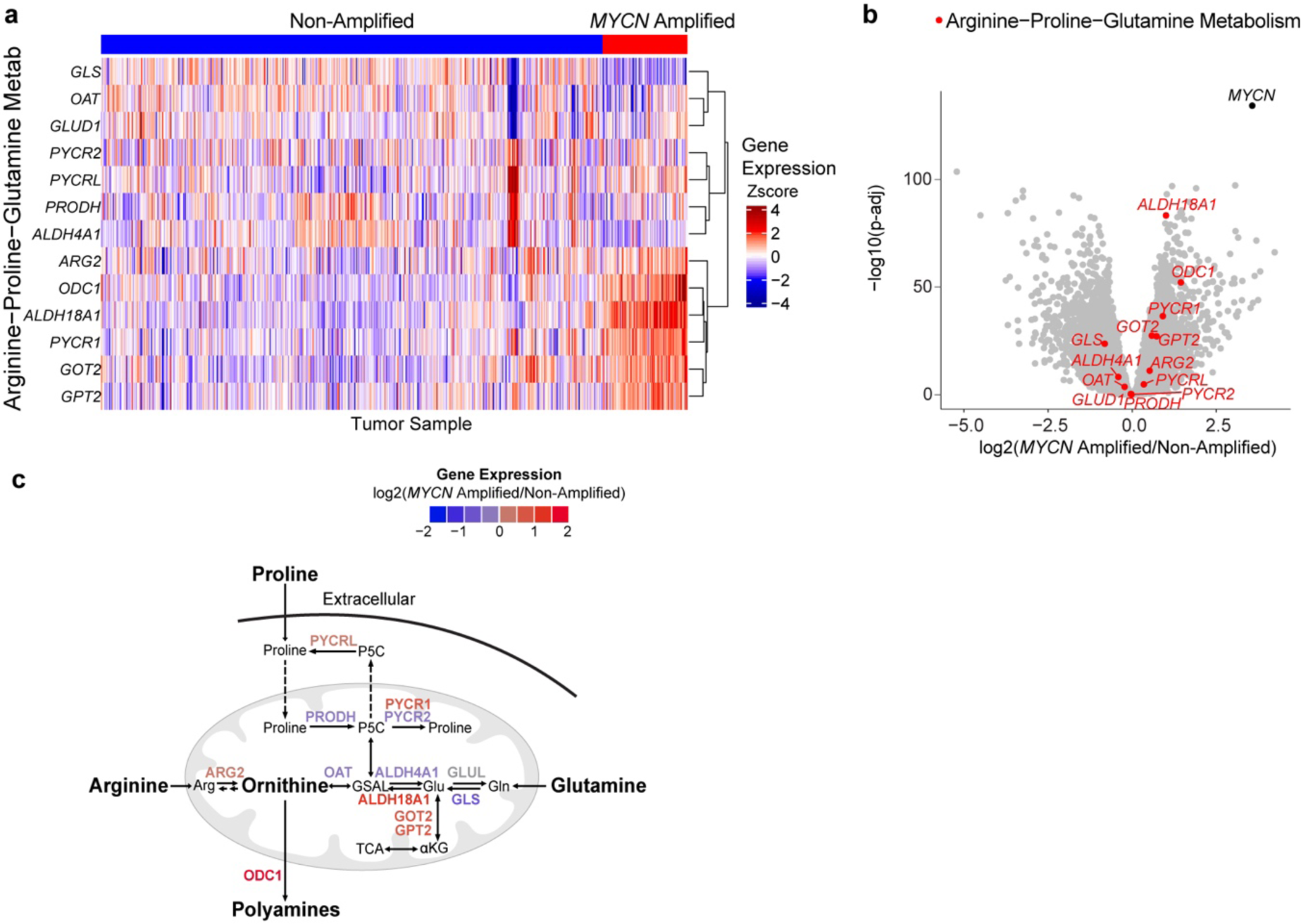
Arginine and proline metabolism expression profiles in primary neuroblastoma patients. a) Heatmap of gene expression from genes related to arginine, proline and glutamine metabolism according to MYCN status. b) Differential gene expression between *MYCN* amplified and non-amplified primary human neuroblastomas. Genes related to arginine, proline and glutamine metabolism denoted in red, *MYCN* in black and all other genes in grey. c) More detailed schematic of gene expression levels of enzymes across arginine, proline and glutamine metabolism with the color of each gene label indicating relative expression (*MYCN* amplified / non-amplified). All graphs display primary neuroblastoma expression data^21^. *MYCN* amplified *n* = 93, Non-amplified *n* = 551. Related to Fig.1e.

**Extended Data Fig. 3:**
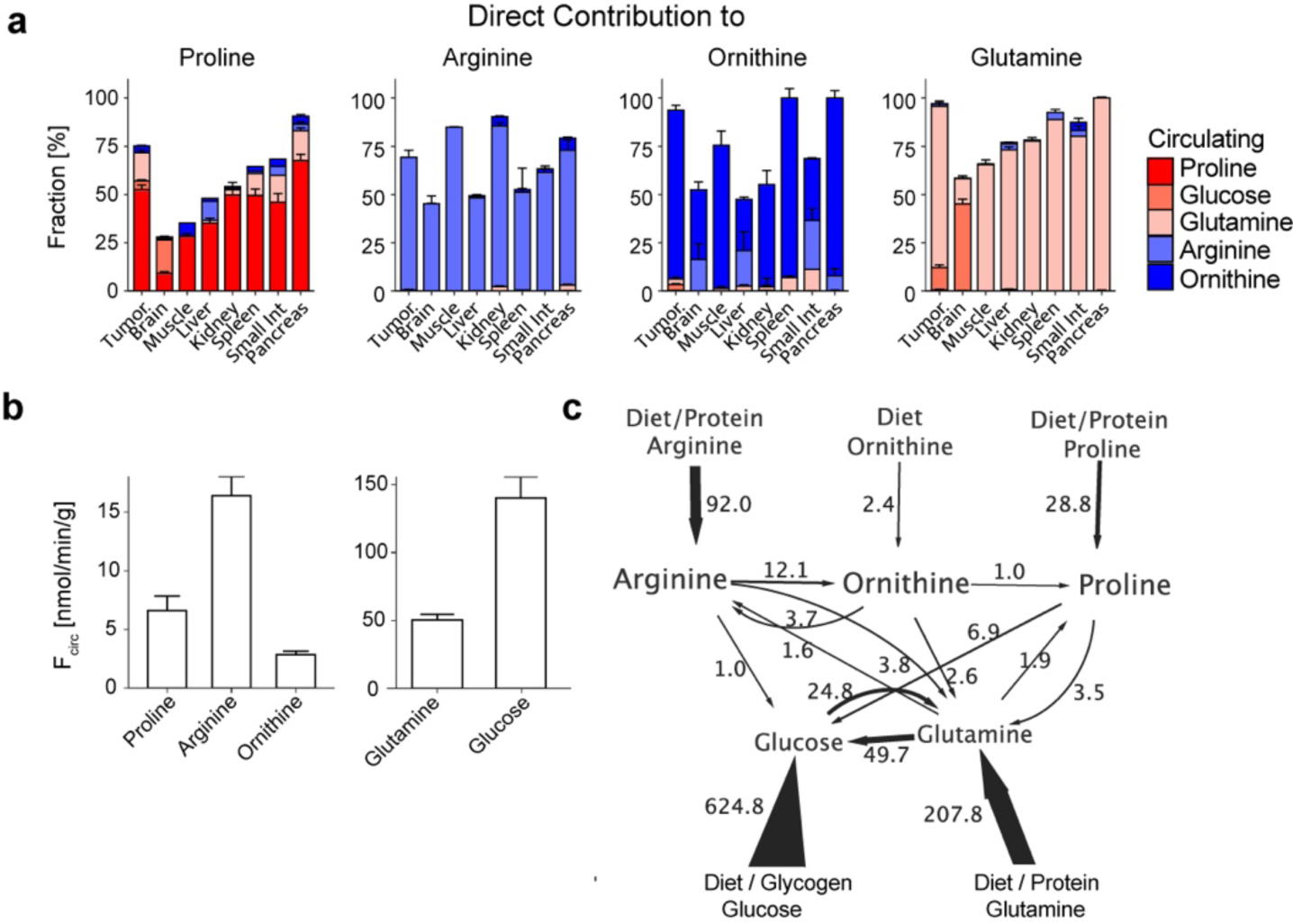
Direct metabolite contributions across organs and turnover flux in the TH-*MYCN* model. a) Direct contributions of circulating nutrients to proline, arginine, ornithine and glutamine across organs and neuroblastoma tumors. Contributions derived from [U-^13^C]-labelled tracer infusions as given in the legend, complementing Figure 1f. Mean ± s.e.m., tumor, pancreas, brain, serum *n* = 4-9; liver, muscle, kidney *n* = 3-7; small int., spleen *n* = 1-3. b) Circulatory serum turnover flux, F_circ_ according to Hui et al.^22^ of proline, arginine, ornithine, glutamine and glucose. Mean ± s.e.m., proline *n* = 4, arginine *n* = 8, ornithine *n* = 6, glutamine *n* = 9, glucose *n* = 4. c) Whole-body flux model of interconversion between sources of circulating proline, ornithine, arginine, glutamine and glucose in nmol C/min/g^22^. Mean fluxes above 1 nmol C/min/g are displayed. Abbreviation: Small int., small intestine.

**Extended Data Fig. 4:**
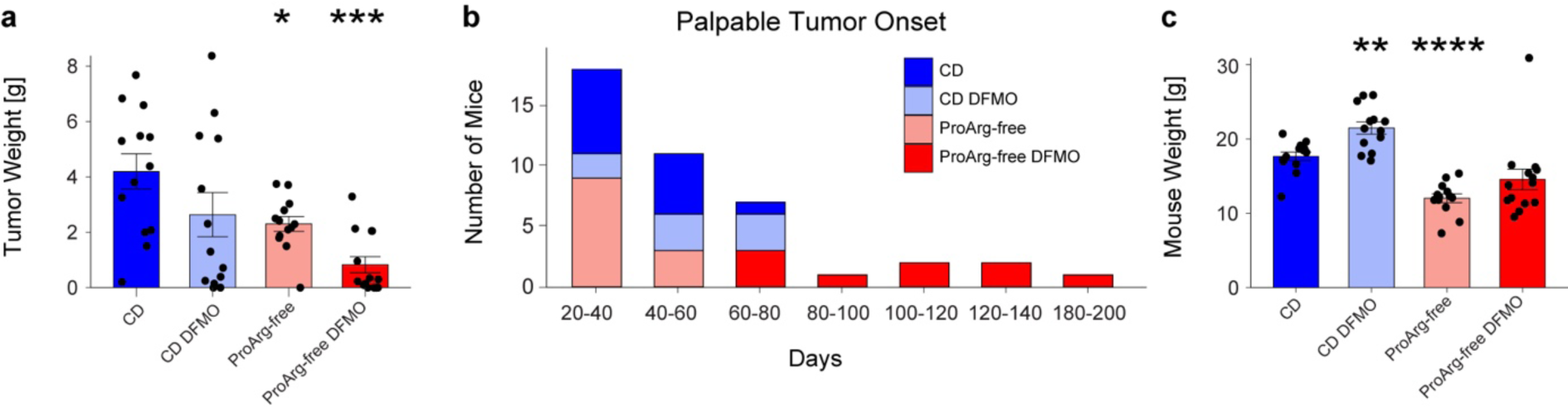
Effect of proline and arginine amino acid depletion combined with DFMO treatment on *MYCN*-driven neuroblastoma *in vivo*. a) Tumor weight at death in grams. b) Delayed tumor growth as shown by time to palpable tumor onset across all treatment arms. c) Mouse weight at death in grams, adjusted for tumor weight. All panels: two-tailed t-test compared to CD, *P < 0.05, **P < 0.01, ***P < 0.001, ****P < 0.0001. CD *n* = 13, CD DFMO *n* = 14, ProArg-free *n* = 13, ProArg-free DFMO, *n* = 14. Abbreviations: CD, control diet; ProArg-free, proline arginine free diet; DFMO, difluoromethylornithine.

**Extended Data Fig. 5:**
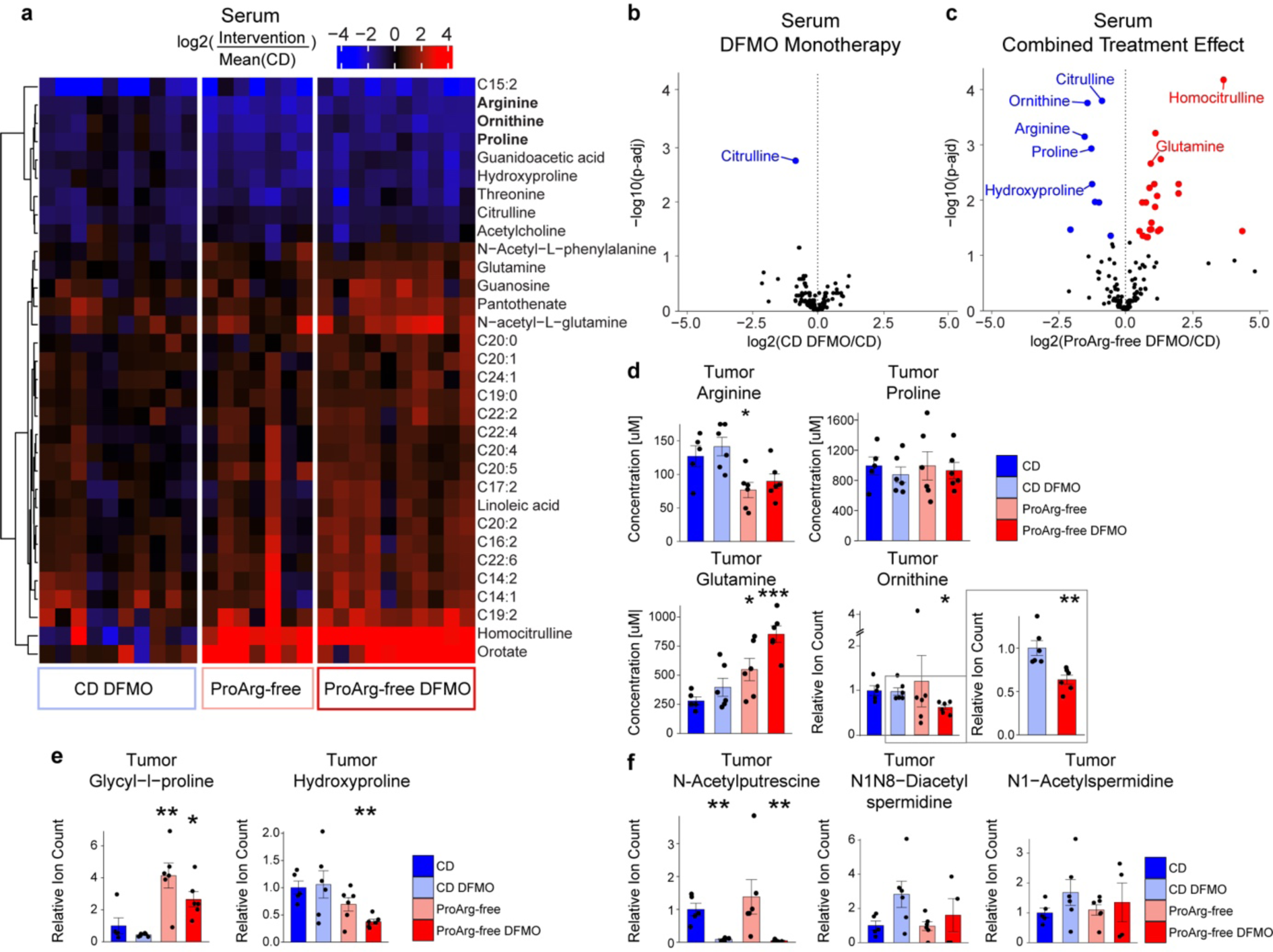
Metabolite profiles of serum and tumors under dietary proline and arginine depletion and / or difluoromethylornithine (DFMO) treatment. a) Heatmap of significantly changed serum metabolites compared to control diet (CD). Metabolites are selected if significantly changed (q < 0.05) in any of the 3 comparisons: Diet effect (ProArg-free vs. CD), DFMO Monotherapy (CD DFMO vs. CD) and Combined Treatment (ProArg-free DFMO vs. CD). Unsupervised clustering performed using Ward’s method. For each intervention, individual mouse are displayed relative to the average level in 7 CD mice. *n* = 75-10. b-c) Differential serum metabolite levels in DFMO Monotherapy and Combined Treatment. Blue highlights metabolites that are significantly depleted (FDR < 0.05) and shades of red up regulated in the treatment group compared to CD. *N* = 7-10. d) Tumor levels of arginine, proline, glutamine and ornithine, compared to CD. *n* = 5-6 e) Tumor levels of proline related metabolites dipeptide glycol-l-proline and hydroxyproline, compared to CD. *n* = 5-6. e) Tumor levels of acetylated polyamines, compared to CD. *n* = 4-6. For A-E metabolomics was performed on prolonged treatment tumors (average sacrifice at 12 weeks in all arms). B-E: *P < 0.05, **P < 0.01, ***P < 0.001, ****P < 0.0001, two-tailed t-test. Mean ± s.e.m.. Technical replicates are averaged for each mouse. Abbreviations: CD, control diet; ProArg-free, proline arginine free diet; DFMO, difluoromethylornithine.

**Extended Data Fig. 6:**
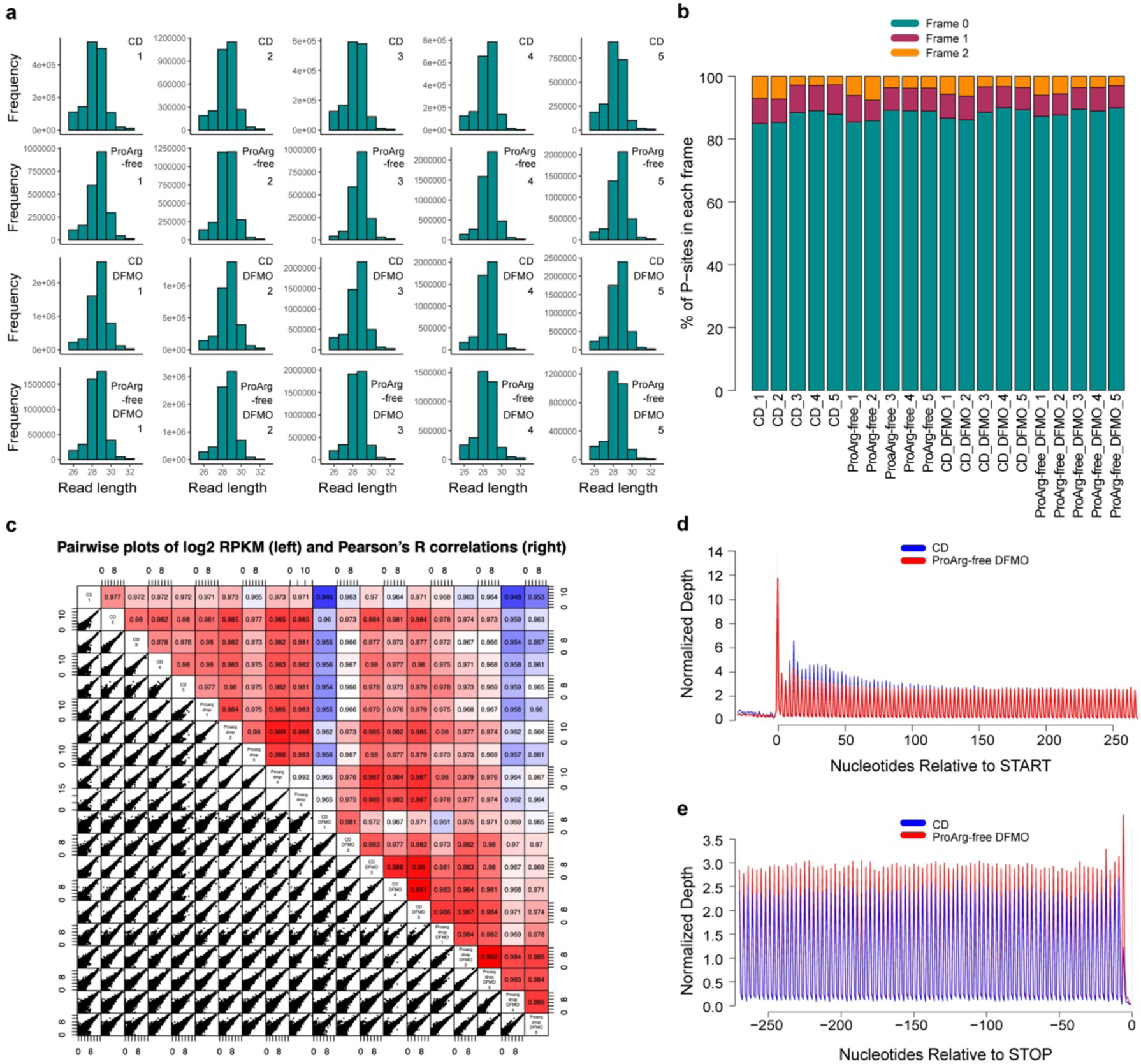
Ribo-Seq quality control. a) Read length frequency distribution of ribosome footprints of each measured sample showing no group specific differences. b) Periodicity by frame for each measured sample showing high frame 0 concordance across the cohort. Mean. c) Pairwise ribosome profile correlation between samples. d-e) Nucleotide resolution ribosome occupancy indicating high periodicity of translation at the transcript 5’ and 3’ comparing CD and ProArg-free DFMO samples. Mean. For A-E: *n* = 5. Abbreviations: CD, control diet; ProArg-free, proline arginine free diet; DFMO, difluoromethylornithine.

**Extended Data Fig. 7:**
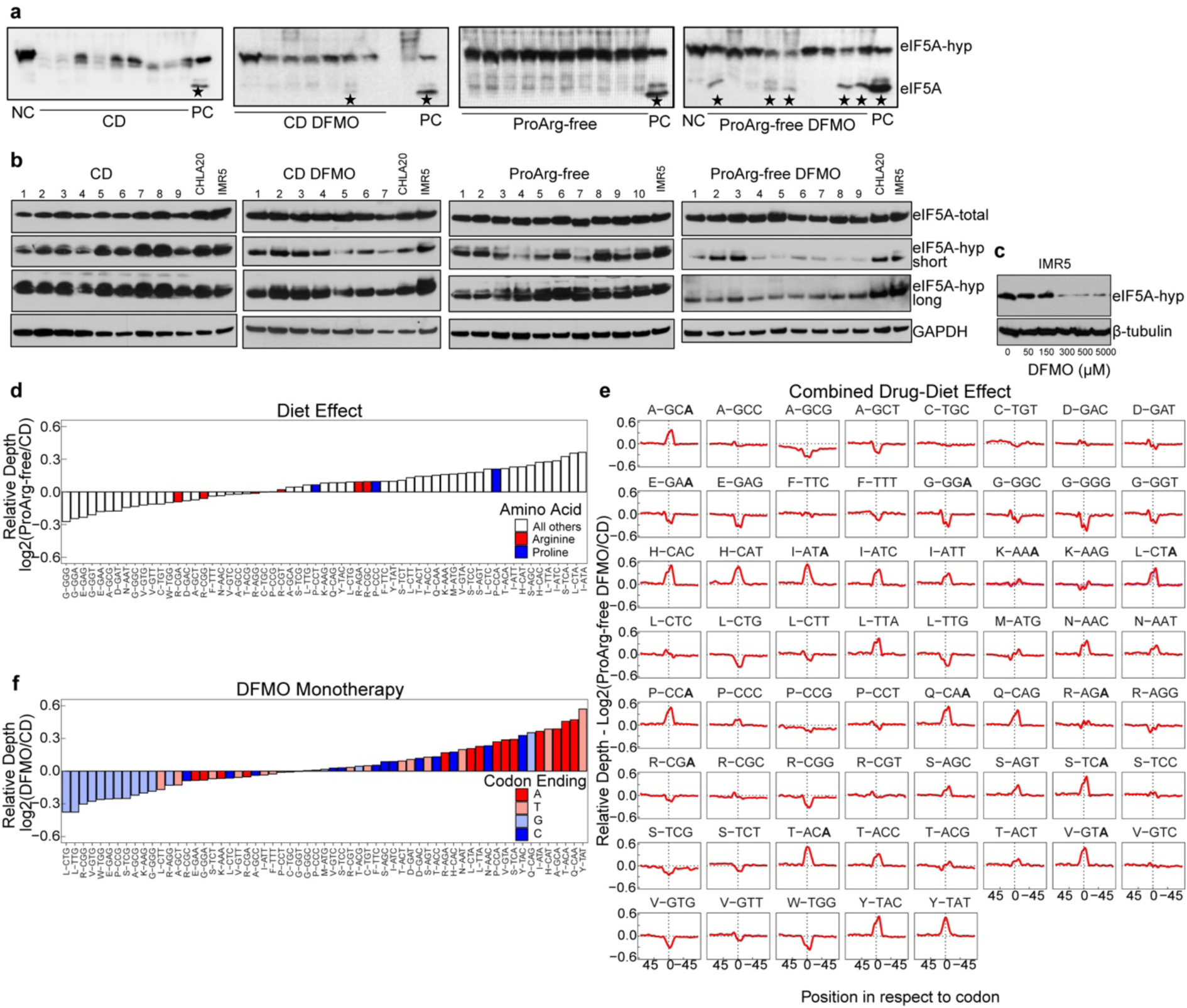
Polyamine depletion causes codon specific translation defects. a) eIF5A hypusination defects detected upon isoelectric focusing followed by immunoblotting. Tumors of ProArg-free and CD DFMO arms, showing defective hypusination in only one CD DFMO tumor and five ProArg-free DFMO tumors. Neuroblastoma cell line IMR5 as negative controls (NC), not treated with DFMO, and positive control (PC), treated for five days with 500 µM of DFMO. Stars denote tumors with non-hypusinated eIF5A (eIF5A) band compared to hypusinated band (eIF5A-hyp). b) eIF5A hypusination defects detected by immunoblotting. Tumors of CD, CD DFMO and ProArg-free show no defective hypusination. ProArg-free DFMO show defective hypusination. IMR5 and CHLA20 were added as negative control for comparison to incomplete hypusination. Hypusinated eIF5A antibody under short (eIF5A-hyp short) and long exposure (eIF5A-hyp long). c) Control neuroblastoma cell line IMR5 treated under increasing DFMO concentrations for 5 days. d) Diet effect (ProArg-free vs. CD) in the ribosome P site is not characterized by ribosome pausing at arginine and proline amino acid codons e) Translation defects are codon specific. Relative ribosome density on all amino acid codons in combined drug-diet treatment (ProArg-free DFMO vs. CD). First letter of name denotes amino acid followed by the encoding codon. f) DFMO treatment effect (CD DFMO vs. CD) is characterized by ribosome pausing depending nucleotide at ending position in ribosome P site. C-G shows mean of *n* = 5. Abbreviations: CD, control diet; ProArg-free, proline arginine free diet; DFMO, difluoromethylornithine,

**Extended Data Fig. 8:**
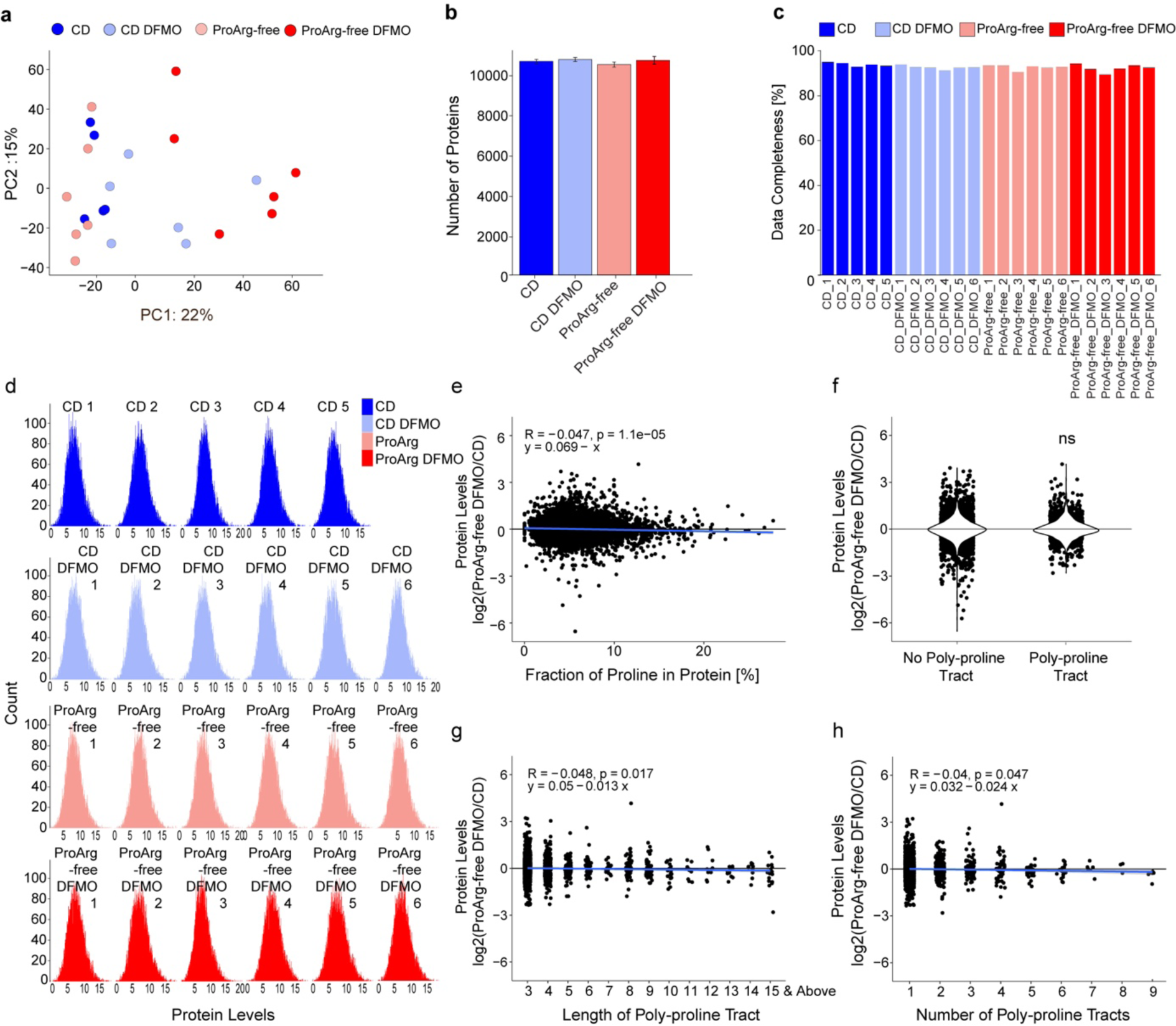
Proteomics quality control and evaluation of protein levels in relation to proline content. a) Global proteomic signatures across all treatment arms analyzed by principal component analysis. b) Number of proteins measured per treatment group. Mean ± s.e.m.. c) Data completeness of each proteomics analytical measurement, with percentage of proteins captured by each sample. d) Distribution per sample of protein levels. e) Correlation between relative protein levels in the combined diet-drug treatment (ProArg-free DFMO vs. CD) and the percentage of prolines from all amino acids of the respective protein. f) Comparison of relative change in protein levels induced by combined diet-drug treatment depending on the presence or absence of at least one poly-proline tract. Poly-proline tracts are defined by having >=3 prolines in a row. g) Relative protein levels in combined diet-drug treatment grouped by the length of poly-proline tract in the protein. h) Change in protein levels in the ProArg-free DFMO group depending on the number of poly-proline tracts in the protein. For a-h: CD DFMO, ProArg-free and ProArg-free DFMO *n* = 6; CD *n* = 5. Abbreviations: CD control diet; ProArg-free proline arginine free diet; DFMO difluoromethylornithine.

**Extended Data Fig. 9:**
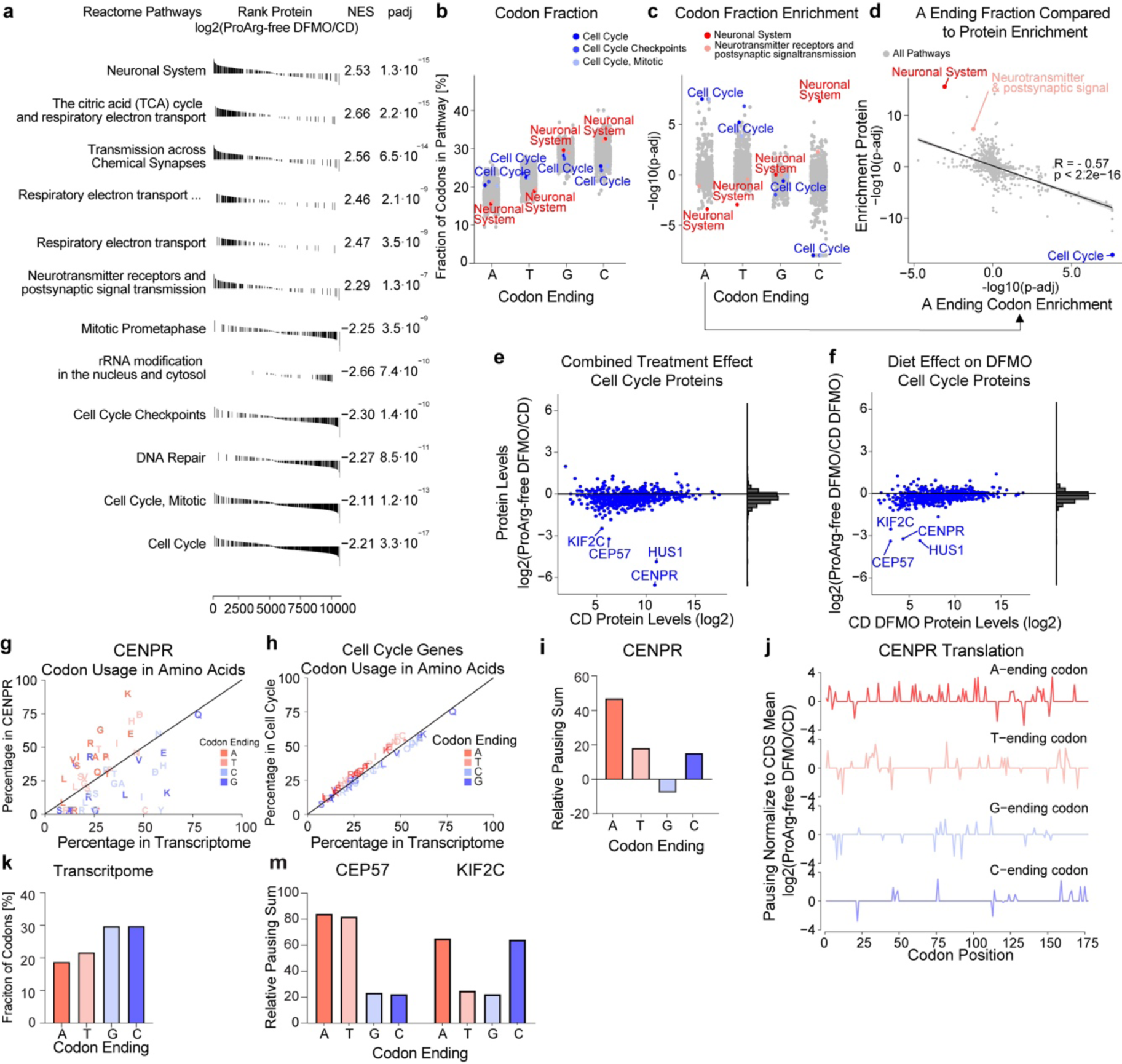
Adenosine-ending codon frequency correlates with translation defects. a) Top down- and upregulated Reactome pathways in the GSEA analysis on the protein level when comparing the diet-drug combination to the control diet (ProArg-free DFMO vs. CD). b) Fraction of codons ending with respective nucleotide across all Reactome pathways (grey), cell cycle related (blue) and neuronal system (red). c) Enrichment analysis based on the A, T, G or C ending codon fraction per gene. Pathways positively enriched (positive p-adj) have higher A, T, G or C ending fractional enrichment whereas negatively enriched pathways have a lower fraction. d) Across Reactome pathways, increasing enrichment of adenosine-ending codons correlates to downregulation on the protein level when comparing combined diet-drug treatment to control. Enrichment was computed using gene set enrichment analysis, where transcripts are ranked by adenosine-ending codon fraction and proteins by fold change on the protein levels. e) Protein intensity across cell cycle, the most downregulated pathway, where the difference between ProArg-free DFMO and CD is evaluated by protein fold-change on the y axis. Fold change distribution on the right side. f) ProArg-free DFMO and CD DFMO highlighting the additive diet effect on the downregulation of cell cycle proteins. Four top-down regulated proteins were identified in both comparisons. Fold change distribution on the right side. g) Relative codon preference comparing the *Itgb3pg* gene (CENPR protein) to the whole transcriptome. Percentages above the line highlight a preferential use for encoding the amino acid in *CENPR* by that specific codon type (red is adenosine ending). h) Relative amino acid codon preference comparing the pathway cell cycle to the whole transcriptome. Marks above the line highlight the preferential utilization of defined codons in the cell cycle genes to encode the same amino acid. i) The relative ribosome pausing sum as the sum of ribosome occupancy ratios of ProArg-free DFMO to CD according to the nucleotide at the ending position in *Itgb3pg* (CENPR protein). Specific dysfunction in decoding of A-ending codons is observed. j) Relative ribosome occupancy across *Itgb3pg* (CENPR protein) reveals a dysfunction in decoding adenosine-ending codons. Occupancy ratios were calculated between ProArg-free DFMO and CD. Summed by the nucleotide at ending position in i). k) Percentage of codons with the respective nucleotide at the ending position in the whole transcriptome. l) Relative pausing sum of *Cep57* and *Kif2c*, where the relative ribosome occupancy ratio between ProArg-free DFMO and CD are summed according to the nucleotide at the codon ending position. Hus1 did not show sufficient coverage. For a, d, e and f: Proteomics ProArg-free DFMO n = 6; CD DFMO n = 6; CD n = 5. For i, j and m: Ribo-Seq *n* = 5; For b-d, the legend for pathways is shared above. Abbreviation: ‘Respiratory electron transport …’ is short for ‘Respiratory electron transport, ATP synthesis by chemiosmotic coupling, and heat production by uncoupling proteins ‘

**Extended Data Fig. 10:**
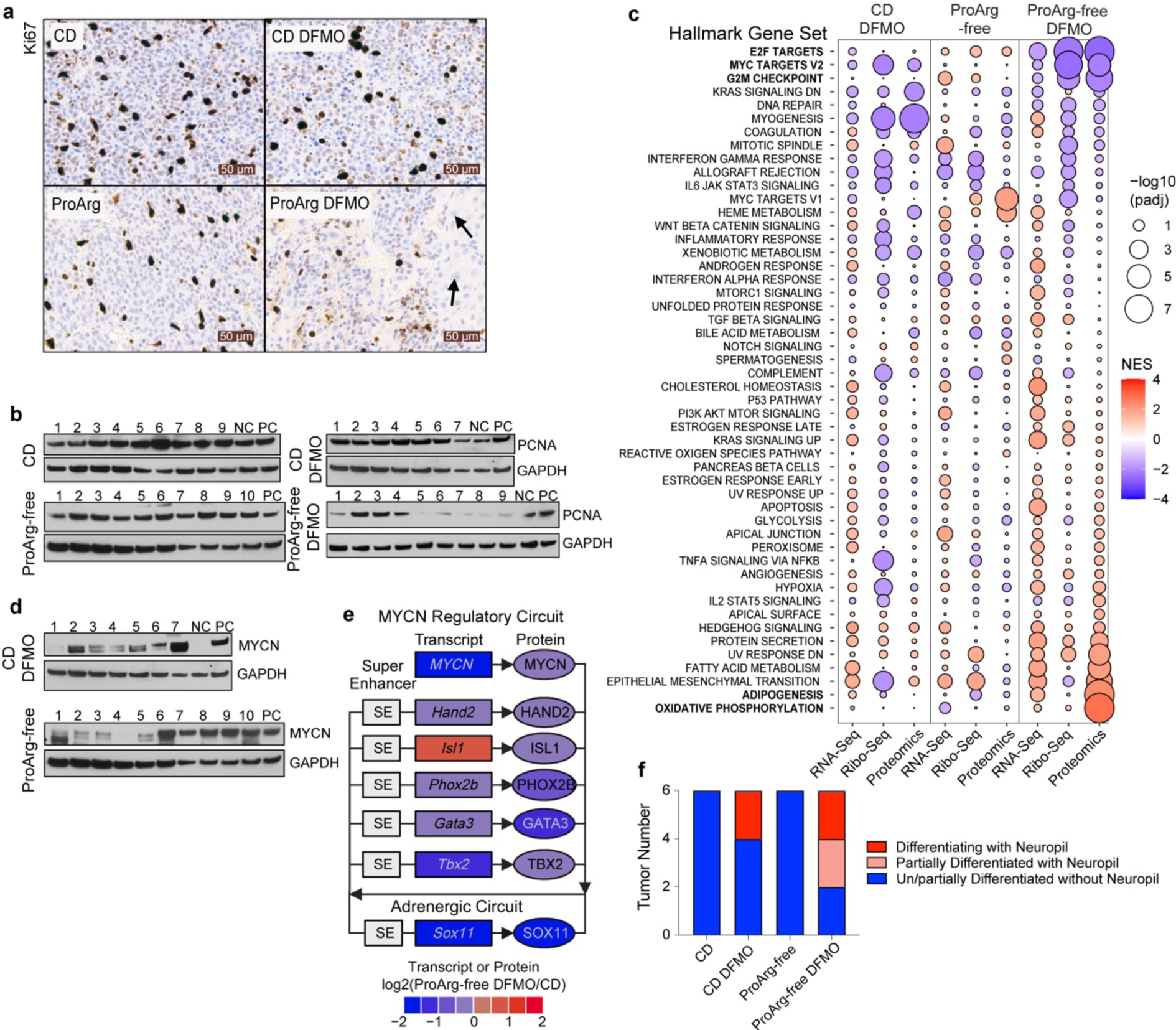
Targeting metabolic dependencies of translation affects hallmarks of cellular regulation. a) Immunohistochemistry showing Ki67 (proliferation marker) of representative TH-*MYCN* tumors. Arrows denote cells in areas with local cytodifferentiation. b) Western blot analysis of PCNA and MYCN in tumors from CD DFMO and ProArg arms. GAPDH is used as loading control. Tumors are identified from 1 to 9. NC denotes negative control CHLA20 neuroblastoma cell line with MYC amplification (not MYCN) and PC denotes positive control IMR5 neuroblastoma cell line (MYCN amplified). c) Hallmark gene set enrichment across omics layers. Displayed is the full gene set tested across each treatment group CD DFMO, ProArg-free and combined ProArg-free DFMO compared to CD. The top 5 significantly changed gene sets on protein level are highlighted in bold. Point size denotes the significance level and the color scale the normalized enrichment score (NES), with red showing enriched in the intervention group (CD DFMO, ProArg-free or ProArg-free DFMO) and blue in CD. d) Western blot analysis of MYCN in tumors from CD DFMO and ProArg-free arms. GAPDH loading control loaded in separate blot (same as PCNA). NC denotes negative control CHLA20 neuroblastoma cell line with MYC amplification (not MYCN) and PC denotes positive control IMR5 neuroblastoma cell line (MYCN amplified). e) Combined diet-drug treatment disrupts the *MYCN* driven super enhancer circuitry on the transcript expression (square), and the protein level (ellipsoid). Similarly, other elements of the core regulatory circuitry are affected^39,40^. f) Blinded histological assessment by pathologist of differentiation status and abundance of neuropil status in tumor sections (linked to Fig.5j). n = 6. For c and e: RNA-Seq: CD DFMO, ProArg-free and ProArg-free DFMO n = 5; CD n = 4, Ribo-Seq: n = 5, Proteomics: CD DFMO, ProArg-free and ProArg-free DFMO n = 6; CD n = 5. Abbreviations: CD, control diet; ProArg-free, proline arginine free diet; DFMO, difluoromethylornithine.

